# Interactome mapping in human excitatory neurons reveals novel risk genes and pathways in Alzheimer’s disease

**DOI:** 10.64898/2026.03.14.711835

**Authors:** Xiaomu Wei, Katie Munechika, Yu Sun, Yuansong Wan, Tianyu Xia, Yuan Hou, Wenqiang Song, Kumar Yugandhar, Yiwen Wang, Se-In Lee, Zhengdong Sha, Yadi Zhou, Weixi Feng, Jingjie Zhu, Yuliang Tang, Wenjie Luo, Feixiong Cheng, Li Gan, Haiyuan Yu

**Affiliations:** Department of Computational Biology, Cornell University, Ithaca, New York, USA; Weill Institute for Cellular and Molecular Biology, Cornell University, Ithaca, New York, USA; Meinig School of Biomedical Engineering, Cornell University, Ithaca, New York, USA; Helen and Robert Appel Alzheimer’s Disease Institute, Feil Family Brain and Mind Research Institute, Weill Cornell Medicine, New York, New York, USA; Department of Genomic Sciences and Systems Biology, Cleveland Clinic Research, Cleveland, Ohio, USA; Cleveland Clinic Genome Center, Cleveland Clinic Research, Cleveland, Ohio, USA; Department of Molecular Medicine, Cleveland Clinic Lerner College of Medicine, Case Western Reserve University, Cleveland, Ohio, USA; Neuroscience Graduate Program, Weill Cornell Medicine, New York, New York, USA

## Abstract

Alzheimer’s disease (AD) is an irreversible neurodegenerative disease defined by its molecular hallmarks - amyloid beta peptide plaques and neurofibrillary Tau tangles. Despite significant progress that has been made in uncovering a large number of genetic risk factors through extensive genomic sequencing and genetic studies, the molecular mechanisms driving AD-associated pathology and cognitive decline remain poorly understood. Therefore, alongside the identification of more risk genes, it is also paramount to study how these genes function and influence each other within the cellular pathways and overall molecular networks in AD-relevant brain cell types. However, current human protein-protein interactome datasets were all generated in either yeast or generic human cell lines. Consequently, many important neuronal interactions, especially neuron-specific ones, have yet been discovered. To address this critical gap, we developed a highly scalable, high-quality interactome mapping pipeline in human excitatory neurons derived from induced pluripotent stem cells (iPSC), and generated a comprehensive, neuron-specific interactome map, named ADNeuronNet, for key AD risk genes. ADNeuronNet consists of 1,767 high-confidence interactions among 1,189 proteins and is the only dataset enriched with neuron-specific genes when compared to known protein interactions, including previous large-scale interactome maps, for the same baits in the literature. Within ADNeuronNet, we identified 1,375 novel interactions, many of which are likely neuron specific. For example, we identified a neuron-specific interactor, RIN2, for major AD risk factor BIN1 and confirmed RIN2’s function in recruiting BIN1 to RAB5 positive early endosomes, a process that has been well-associated with AD etiology. Additionally, we performed quantitative interaction perturbation analyses on AD risk genes with AD-associated mutations or isoforms and identified significant changes in 99 protein interactions among 11 different protein variants. Finally, we found that subunits from the anaphase-promoting complex/cyclosome (APC/C), another novel BIN1 interactors identified by ADNeuronNet, mediated modulation of Tau-aggregation in neurons via regulation of APOE expression, uncovering a previously unrecognized BIN1-APC/C-APOE regulatory axis in AD pathobiology. In summary, these findings illustrate how our neuron-specific ADNeuronNet can be leveraged to uncover new risk gene candidates and cellular pathways that help advance our understanding of molecular mechanisms underlying AD etiology.

## Introduction

Alzheimer’s Disease (AD) is an irreversible neurodegenerative disease accounting for sixty to eighty percent of all dementia cases. An estimated 7.2 million individuals in the United States are living with AD currently, a number projected to double by 2050^1^. Pathological amyloid beta (Aβ) plaques and phosphorylated Tau neurofibrillary tangles define the molecular hallmarks of AD. Yet mechanisms of cognitive decline remain elusive^2,3^. Large scale genomic efforts, including the AD Sequencing Project^4^ and genome wide association studies (GWAS), have transformed our understanding of AD genetics in the last decade, identifying ∼100 susceptibility loci that contribute to late onset sporadic AD, which represents more than ninety five percent of all cases^5–7^.

Despite this remarkable progress, the biological mechanisms by which AD risk genes initiate and drive disease remain poorly defined. Most risk variants confer modest effects, act in a polygenic and context dependent manner, and exhibit pleiotropy across cell types and disease stages. As a result, genetic association alone has proven insufficient to explain how risk alleles converge on pathogenic pathways. Bridging this gap requires moving beyond gene lists toward a mechanistic framework that captures how disease associated proteins function within cellular systems.

Protein function is dependent on interactions with other proteins^8,9^. Recent studies have shown that disruption of specific protein-protein interactions underlies the molecular mechanism of many disease genes^10–13^. Therefore, integration of multi-omics data within the framework of the protein-protein interaction network (often called the interactome) can help better dissect the pleiotropy of disease genes and complex genotype-to-phenotype relationships^10,14,15^.

The utilization of protein-protein interaction information is increasingly recognized as an essential component for elucidating the cellular machinery of AD. Interactome mapping efforts focused on key AD proteins, such as Tau^16–18^, APP^19^, and Aβ^20^, have recently been carried out. Tau interactomes, which have been studied in patient cortical tissue^18^, NFT samples^16^, and iPSC-derived neurons^17^ have revealed intriguing connections relating Tau with proteasome pathways, mitochondrial proteins, and synaptic function via interactions with SNARE complexes. Other studies have focused on validating Aβ interactors in human cortex tissues^20^ or the interactome of APP in transgenic mouse models^19^, which emphasized the involvement of APP with synaptic vesicle processes. Although significant insights were gained from these interactomes regarding the functional roles of these proteins, each study focused on only one bait protein at a time. This limited scope does not allow the ability to capture connecting interactions and potentially converging pathways among a large number of different risk proteins. Other efforts by CCSB^21^, BioPlex^22^, and OpenCell^23^ Consortia have worked to systematically map the global protein-protein interactome network involving thousands of human proteins for over a decade using high-throughput yeast two-hybrid (Y2H)^21^ or large-scale immunoprecipitation-followed-by-mass-spectrometry (IP-MS)^22,23^. These mapping efforts and the detected interactions are extremely valuable, but most of the currently-known interactions for human proteins are detected in yeast cells^21^ or generic human cell lines (mainly HEK293T^22,23^ and HCT116^22^). As a result, these datasets may be missing information regarding brain-cell-specific interactions that can play key roles in brain functions and AD pathogenesis. In order to gain a comprehensive view of the biological processes and reflect the complex interconnectedness of disease-associated molecular pathways, it is necessary to build a global AD interactome, containing all risk proteins, in a clinically-relevant brain cell-type.

Glutamatergic excitatory neurons constitute the majority of neurons in the central nervous system and represent a primary site of selective vulnerability in AD^24^. These cells are early sites of Tau accumulation and key drivers of network dysfunction in AD. Spatial transcriptomics shows that Tau positive excitatory neurons in both primary age related tauopathy and AD share a conserved, cell intrinsic transcriptional program enriched for synaptic and calcium homeostasis pathways despite global synaptic downregulation. This signature reflects a self-reinforcing cascade in which disrupted glutamate signaling, calcium dysregulation, and Tau accumulation converge to impair synaptic structure and excitability^25^. Mapping how AD risk genes and their protein interactors function within glutamatergic neurons could link genetic risk with cell type specific mechanisms of synaptic remodeling, calcium imbalance, and Tau driven neurodegeneration.

Towards this goal, we developed a highly scalable high-quality interactome mapping pipeline in human iPSC-derived excitatory neurons using quantitative IP-MS, and produced the first comprehensive human neuron specific protein-protein interactome network for 57 AD risk proteins, named ADNeuronNet. Using an optimized data independent acquisition, label free quantification (DIA-LFQ) workflow, we identified 1,767 high-confidence interactions among 1,189 proteins in human excitatory neurons, 1,375 of which are completely novel, many are likely neuron-specific. In fact, compared with known protein interaction datasets in the literature, our new interactome is the only one enriched with neuron-specific genes, highlighting the necessity of studying genes important for neuronal functions in human neurons and confirming the validity of our overall approach. Notably, we found previously unreported interactions of BIN1, a major genetic risk factor for AD, with RIN2 and APC/C complex. Specifically, RIN2 is not expressed in either HEK293T or HCT116 cells and therefore was unable to be identified by previous interactome mapping efforts. Yet, RIN2 is highly expressed in neurons and across key brain regions, and our interactome and subsequent cell biology results implicate its functional roles in AD etiology. Furthermore, the novel BIN1-APC/C interaction and our follow-up multifaceted functional studies reveal a previously unrecognized BIN1-APC/C-APOE regulatory axis in human neurons and demonstrate that loss of APC/C function promotes Tau aggregation through aberrant induction of neuronal APOE, highlighting how our neuron-specific interactome, ADNeuronNet, can be utilized to discover new candidate genetic drivers that can impact AD pathobiology, some of which can potentially be effective therapeutic targets.

## Results

### Comprehensive interactome mapping for AD risk genes in human iPSC-derived neurons

To build a list of the most significant AD risk genes, we included all 86 high-confidence protective or causal genes with genetic and functional evidence compiled by the Alzheimer’s Disease Sequencing Project (ADSP)^4^, as well as all 114 genes from the 3 latest AD GWAS studies^5,26,27^ (see **Methods**). These studies prioritized 89 risk genes from GWAS loci through comprehensive and advanced analyses. For comprehensiveness, all nearest genes to all significant lead SNPs were also included (25 additional genes)^5,26,27^. In total, we compiled a list of 137 unique AD-associated genes with strong genetic support.

Next, we checked the neuronal expression of these genes and found 97 genes to be expressed in neurons based on transcriptome data (see **Methods**)^28^. We successfully cloned 57 of the 97 neuron-expressed AD risk genes into our lenti-viral expression vector with a GFP tag for downstream experiments (**Fig. 1a**). The 40 failed genes have significantly longer cDNAs (**Fig. S1a**), which made the cloning much more challenging. Additionally, we also generate clones for 11 key AD-related isoforms and patient derived mutations. Specifically, Bridging Integrator 1 (BIN1) is the second most significant AD risk gene following APOE^5^. The BIN1 gene contains 20 exons that can be spliced into tissue-specific isoforms, with the largest isoform, isoform 1 (BIN1v1), predominantly expressed in neurons^29^. However, previous human interactome mapping efforts^21–23^ exclusively focused on the smaller, ubiquitously expressed isoform 9 (BIN1v9). Here, we included both BIN1 isoforms in our neuron-specific interactome map.

**Figure 1.**
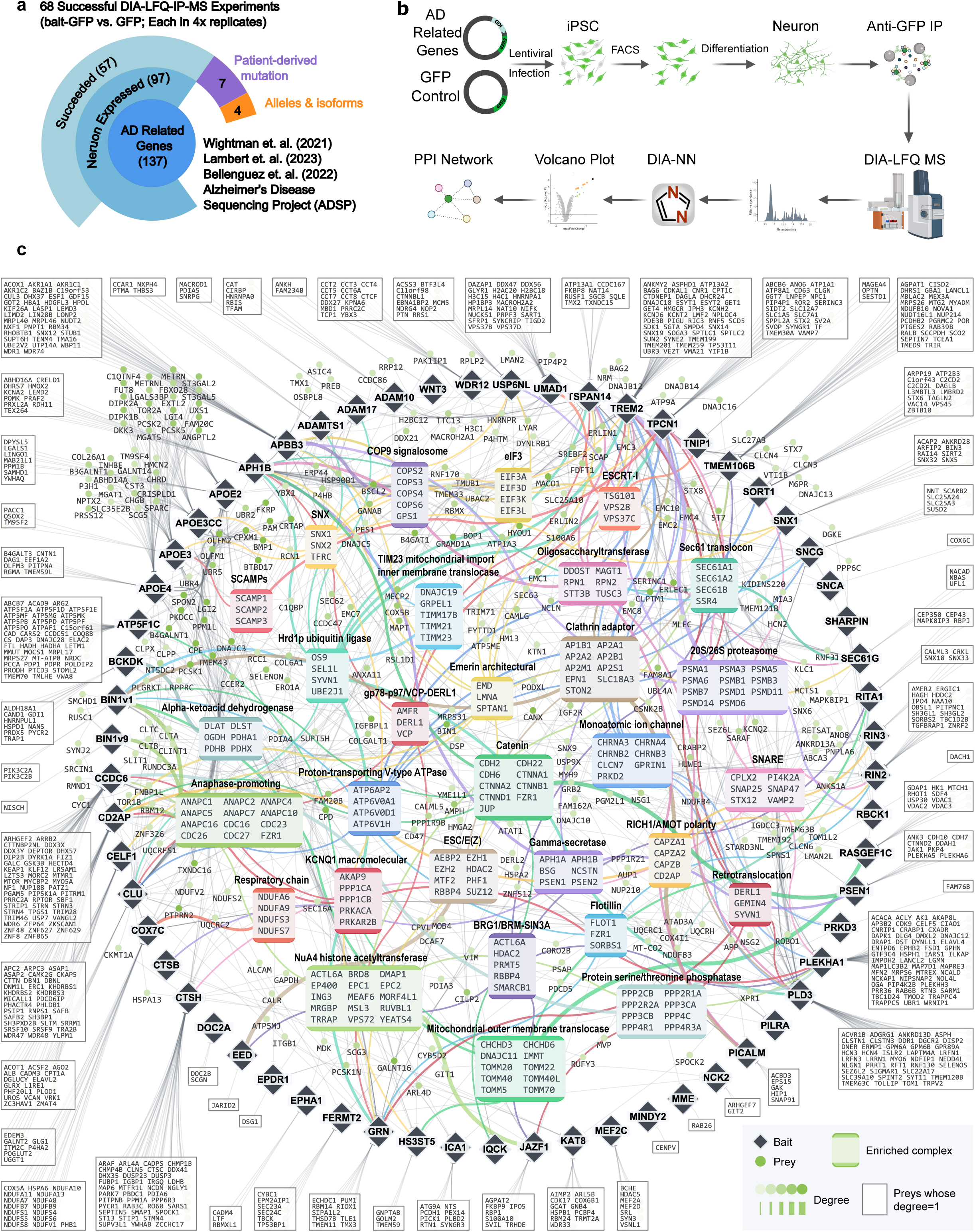
Systematic construction of ADNeuronNet, the neuron-specific interactome for AD risk proteins. **(a)** Diagram representing the numbers of total, neuron expressed, and successfully analyzed AD risk genes. Of the original 137 risk gene list compiled from the literature (see **Methods**), 97 are neuron-expressed and IP-MS was successfully performed for 57 of them, with the addition of 7 AD-associated mutations and 4 isoforms. **(b)** Workflow to perform IP-MS experiments in iPSC-derived neurons. Lentiviral transduction followed by fluorescence activated cell-sorting (FACS) is used to establish iPSC lines stably expressing each AD risk gene and respective controls. After differentiation into neurons, immunoprecipitation is performed to isolate the protein bait and its interactors, followed by mass spectrometry analysis via a DIA-LFQ pipeline. Created in BioRender. **(c)** Network visualization of ADNeuronNet. The black diamond nodes represent AD-associated bait proteins; the circular nodes represent interacting prey proteins; the colorful squares represent enriched protein complexes. Edge thickness indicates the number of interactors from each bait enriched in a given complex. Gray edges show protein-protein interactions within the network.

To establish a scalable iPSC-differentiation based high-throughput IP-MS pipeline, we utilized a doxycycline-inducible neurogenin 2 (NGN2) human iPSC line (i3N) that differentiates into functional glutamatergic cortical neurons (i3Neurons)^30^. This cell line offers a simplified and rapid, yet robust differentiation process making it suitable for high-throughput applications^31^. Through a careful optimization process, we determined that lentiviral transduction of the iPSCs followed by fluorescence-activated cell sorting (FACS) of GFP-positive cells to establish stable iPSC lines expressing each AD risk protein was necessary to obtain a sufficient amount of material for successful IP-MS (**Fig. S1b**). After differentiation, the bait-expressing neurons and control neurons expressing the empty GFP vector were subjected to immunoprecipitation performed in quadruplicates, then analyzed via mass-spectrometry (**Fig 1b**).

We have used TMT-based Data-Dependent-Acquisition (DDA-TMT) workflow for our past interactome mapping effort^32^. However, with recent development of Data-Independent-Acquisition (DIA) methods, new label free quantification (DIA-LFQ) workflows have been shown to be even more powerful, providing much increased sensitivity and coverage^33,34^. We thus optimized a customized DIA-LFQ workflow on our Bruker timsTOF HT mass spectrometer. Using a benchmarking dataset containing yeast proteome spiked into human proteome at various ratios, we compared the performances of the DDA-TMT and DIA-LFQ methods. We found that our DIA-LFQ workflow significantly outperforms our previous DDA-TMT workflow with >3× proteome coverage with slightly better quantitative performance while requiring <50% of input materials (**Fig. S2**).

Using our optimized iPSC-derived neuron based high-throughput DIA-LFQ IP-MS pipeline, we successfully carried out IP-MS experiments for all 57 AD-related bait proteins that we cloned. We have also developed a customized analysis pipeline to process all of our DIA-LFQ IP-MS experiments (see **Methods**). With careful normalization and stringent protein abundance and detected precursor count filters, we implemented a robust empirical Bayes moderated statistical framework to detect high-quality interactors for each bait. Overall, we detected 1,769 interactions in human iPSC-derived excitatory neurons, establishing the first large-scale neuron-specific interactome for AD-associated risk proteins, which we named ADNeuronNet (**Fig. 1c**).

To understand the functional organization of the human AD protein interactome, we performed comprehensive protein complex enrichment analysis on interactors. The interactome network revealed enrichment in more than 30 functional protein complexes with AD-relevant biological functions (**Fig. 1c**). Key enriched protein complexes included the anaphase-promoting complex (APC/C), involved in cell cycle regulation and synaptic plasticity; protein serine/threonine phosphatases, key regulators of multiple cellular processes including lysosome organization and protein localization; and mitochondrial outer membrane translocase complex, essential for mitochondrial protein import and mitochondrial function. Notably, multiple baits converged on shared protein complexes, such as BIN1v1 and BIN1v9 sharing the APC/C complex, APH1B and PSEN1 sharing the catenin complex. Some baits also showed specific complex interactions, such as BIN1v1 interacting with the mitochondrial outer membrane translocase complex (e.g., TOMM22, TOMM40 and TOMM40L), and APOE3 interacting with the SCAMP complex. This systematic protein complex mapping demonstrates that AD risk genes interact with a set of core cellular machineries, providing potential mechanistic insights for AD pathogenesis.

### High-quality ADNeuronNet reveals neuron-specific and disease-associated interactors

To ensure the quality of ADNeuronNet, we compared our identified protein interactions with known interactions from the literature, as well as with other large-scale interactomes, e.g., BioPlex (in HEK293T and HCT116 cells)^22^ and OpenCell (in HEK293T cells)^23^ (**Fig. 2a**). We observed significant enrichment of ADNeuronNet interactions with all three datasets, confirming the high quality of our neuron-specific interactome mapping pipeline.

**Figure 2.**
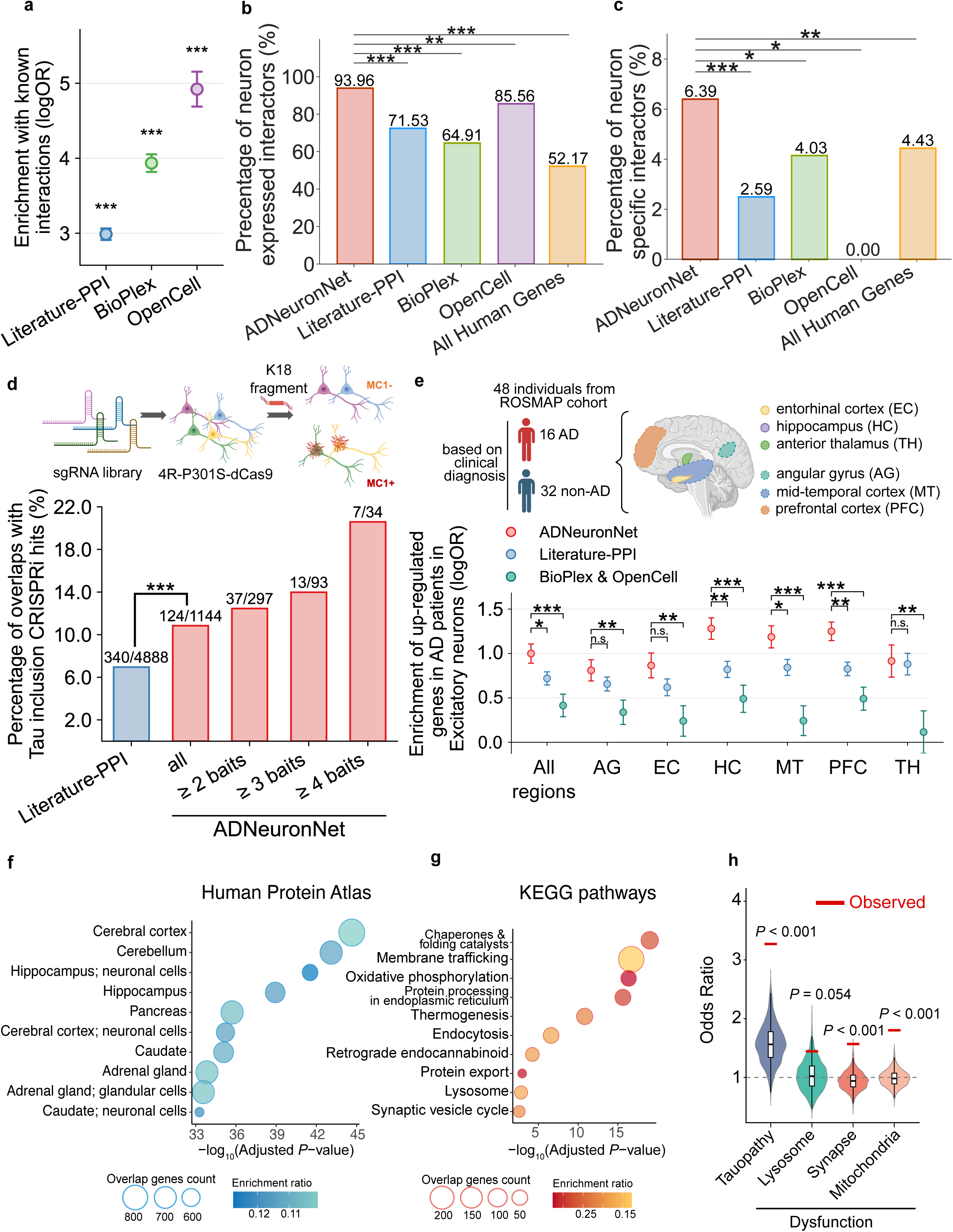
ADNeuronNet is of high quality and relevant to AD pathobiology. **(a)** Enrichment of known protein interactions detected in ADNeuronNet. Known interactions in published human interactome datasets include all known protein-protein interactions previously reported in the literature (Literature-PPI) and other major large-scale human interactome efforts (BioPlex and OpenCell). **(b)** Percentage of neuron-expressed interactors in ADNeuronNet and other published human interactome datasets compared to all human proteins. **(c)** Percentage of neuron-specific interactors in ADNeuronNet and other published human interactome datasets compared to all human proteins. **(d)** Overlap of ADNeuronNet interactors with Tau inclusion-associated CRISPRi hits. The CRISPRi screening strategy utilized MC1 immunoreactivity to endogenous Tau inclusions produced in a 4R-P301S-dCas9 iPSC-derived neuron line as described in Bravo et al ^31^. Interactors in ADNeuronNet are further categorized by the number of AD risk baits that each was found to interact with. Difference with the literature-based interactors was assessed by two-proportion Z-test. Schematic created in BioRender. **(e)** Enrichment of up-regulated genes in clinically diagnosed AD dementia patients in excitatory neurons. Differentially expressed genes were derived from snRNA-seq analysis of ROSMAP samples (16 AD, 32 non-AD). Enrichment was calculated as log odds ratio using Fisher’s exact test and compared to interactors in literature protein interactions and combined BioPlex & OpenCell interactomes. Pairwise differences were assessed using Z-test based on the standard errors of the log odds ratios. Analysis schematic created in BioRender. **(f-g)** Enrichment analysis of interactors: **(f)** Human Protein Atlas (HPA) cell type/tissue enrichment. **(g)** KEGG pathway enrichment in AD-relevant processes. Bubble size indicates overlap gene count; color intensity indicates enrichment ratio. **(h)** Enrichment of proteins associated with AD endophenotypes when compared with randomized PPI background. The red lines show observed values for ADNeuronNet interactors; violin plots show random distributions. *p* indicates significant enrichment over random expectation. Statistical significance is indicated by asterisks (n.s., not significant; \**p* < 0.05; \*\**p* < 0.01; \*\*\**p* < 0.001)

Next, we examined the neuronal expression levels of interactors detected in our ADNeuronNet, and observed high enrichment of neuron-expressed genes within ADNeuronNet (∼95%), significantly higher than the 3 literature interaction datasets. More importantly, compared with the published datasets (most were detected in HEK293T cells and some in HCT116 cells), our ADNeuronNet is the only dataset with significant enrichment of genes specifically expressed in human neurons, but not in either HEK293T or HCT116 cells (**Fig 2b,c; Methods**). These results confirm the neuron specificity of ADNeuronNet, which will help better understand protein functions within neurons and their potential involvement in AD.

In order to investigate the association of ADNeuronNet with proteins known to be associated with AD pathology, we evaluated the overlap of ADNeuronNet interactors with the most significant hits from a CRISPRi screen that determined genes with strong correlations to Tau pathobiology^31^ (**Fig. 2d**). We found that ADNeuronNet interactors had a significantly higher overlap with the CRISPRi hits compared to those already published in the literature. Furthermore, this overlap was more pronounced for proteins that interact with multiple baits (i.e., known AD risk proteins) within ADNeuronNet.

Furthermore, using the latest ROSMAP snRNA-seq data^35^ covering 1.3 million single nuclei from 283 post-mortem human brain samples across 48 individuals with and without AD dementia based on the clinical diagnosis, we find that interactors in our neuron-specific interactome have the highest enrichment of up-regulated genes in AD patients in excitatory neurons across all brain regions relevant to AD (**Fig. 2e**). A similar enrichment pattern was observed when AD status was defined using pathological diagnosis (**Fig. S3a**). In addition, when enrichment was evaluated across different brain cell types, the strongest enrichment was observed in excitatory and inhibitory neurons, which is consistent with the neuron-specific nature of ADNeuronNet (**Fig S3b,c**). These results confirm that our neuron interactome map identifies proteins relevant to neuronal functions and potentially AD etiology.

To explore the biological and functional relevance of ADNeuronNet, we first conducted cell-type-specific enrichment analysis based on the expression levels reported in the Human Protein Atlas^28^. The interactors in ADNeuronNet are significantly enriched in brain-relevant cell types and tissues (**Fig. 2f**). The top enriched brain regions include cerebral cortex, cerebellum, and hippocampus; and the most enriched cell type is neuronal cells within these regions, demonstrating that our neuron-derived interactome map, ADNeuronNet, captures biologically relevant proteins expressed in AD-affected brain regions. Furthermore, KEGG pathway enrichment analysis showed that our interactors enriched on key AD-relevant pathways (**Fig. 2g**), including membrane trafficking, oxidative phosphorylation, protein processing in endoplasmic reticulum, endocytosis, lysosome, and synaptic vesicle cycle. Finally, we examined the enrichment of proteins associated with AD endophenotypes^36,37^. We found that interactors in our ADNeuronNet show significant enrichment in multiple AD-associated endophenotpyes, including tauopathy and dysfunction in lysosome, synapse, and mitochondria.(**Fig. 2h**). These comprehensive systems biology analyses confirm that our neuron-specific ADNeuronNet identifies biologically relevant pathways and brain region-specific neuronal proteins with direct implications for AD pathobiology.

To further demonstrate the quality and scope of ADNeuronNet, we highlight several key protein complexes detected, including the gamma secretase, SNARE, and mitochondrial respiratory chain complexes, that we were able to recapitulate almost completely through our IP-MS experiments (**Fig. 3a-c**). Notably, a few of these interactors are neuron-specific, reflecting our pipeline’s neuron-centric nature. All of these complexes play crucial roles in maintaining neuronal health and have been reported to undergo some type of dysfunction during AD pathogenesis. Gamma secretase, responsible for mediating cleavage of amyloid precursor protein (APP) into Aβ peptides, is especially relevant for AD, as its abnormal activity directly contributes to the accumulation of neurotoxic, aggregation-prone Aβ^38^. For PSEN1, a subunit of the gamma secretase complex, we observe known connections to cadherin and catenin proteins responsible for cell adhesion regulation (**Fig. 3a**)^39,40^. Regarding synaptic function, we observe 10 baits are associated with soluble N-ethylmaleimide-sensitive factor attachment protein receptor (SNARE) proteins (**Fig. 3b**), which facilitate synaptic vesicle exocytosis involved in neurotransmitter release. In an AD context, impaired function or assembly of SNARE proteins can lead to synaptic dysfunction, a phenomenon occurring in the early stages of AD and highly correlated with the severity of cognitive decline^41^. Previous findings have noted that SNAP25, VAMP2, and SYN1 have reduced levels in AD brain samples compared to healthy controls^42^. Furthermore, other studies have shown that intracellular Aβ oligomers can directly bind to SYN1, impairing SNARE assembly formation and inhibiting SNARE-mediated exocytosis^43^. The third complex group we identified in ADNeuronNet are 3 of the 5 main complexes (**Fig. 3c**) making up the mitochondrial electron transport chain, anchored by the cytochrome oxidase c subunit (COX7C) bait and interacting with synuclein protein baits SNCA and SNCG. SNCA’s co-localization and interplay with mitochondrial Complex IV has also been previously reported^44^, though much less is known about SNCG. Tau (MAPT) was also observed interacting with COX7C. This connection between Tau and mitochondrial proteins has also been documented in our previous Tau interactome study^17^, further establishing the association of pathological Tau in AD-related mitochondrial dysfunction.

**Figure 3.**
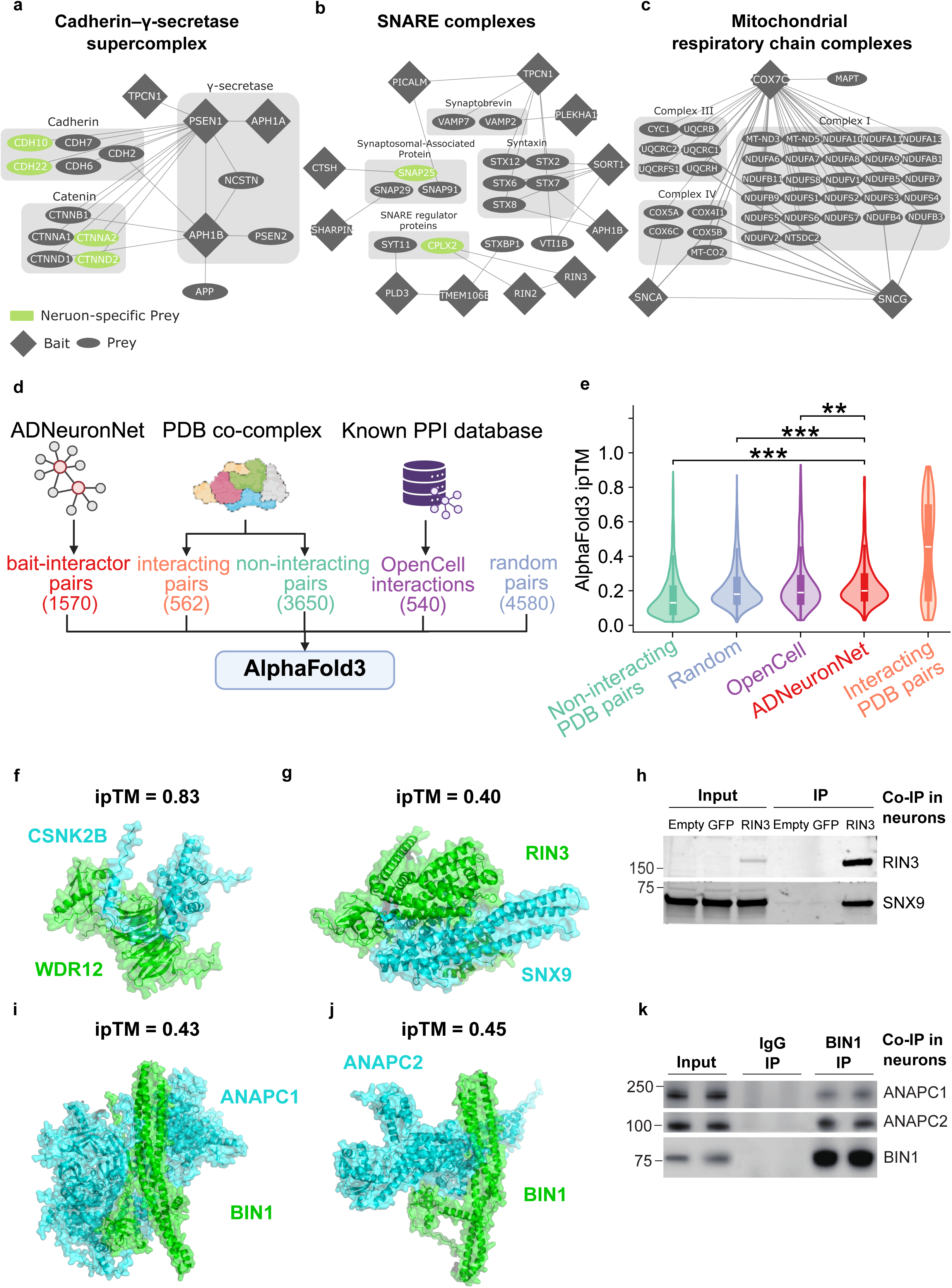
Structural modeling of ADNeuronNet interactions using AlphaFold3. **(a-c)** Network schematics of protein complexes identified within ADNeuronNet, including the cadherin–γ-secretase supercomplex **(a)**, mitochondrial respiratory chain complexes **(b)**, and SNARE complexes **(c)**. AD risk bait proteins are shown as diamonds and interacting proteins as ovals; neuron-specific interactors are highlighted in green. **(d)** Workflow for AlphaFold3-based structural modeling of ADNeuronNet interactions, OpenCell interactions, curated protein–protein co-complexes with available PDB structures, and random protein pairs. Created in BioRender. **(e)** Distribution of AlphaFold3 inter-protein TM-score (ipTM) values across different interaction datasets indicates high quality of ADNeuronNet interactions. Pairwise differences were assessed using Mann-Whitney U test. **(f–g)** Representative AlphaFold3 structural models of protein–protein interactions lacking experimentally resolved structures. **(f)** Previously reported interaction between WDR12 and CSNK2B. **(g)** Novel interaction identified in ADNeuronNet between RIN3 and SNX9. **(h)** Co-immunoprecipitation with anti-GFP beads of the RIN3–SNX9 interaction in human iPSC-derived neurons. ‘Empty’ lanes represent uninfected neurons, ‘GFP’ lanes represent neurons expressing GFP alone, and ‘RIN3’ lanes represent neurons expressing GFP-tagged RIN3. **(i–j)** Representative AlphaFold3 models of previously uncharacterized BIN1 interactions identified in ADNeuronNet. **(i)** BIN1–ANAPC1 and **(j)** BIN1–ANAPC2, both components of the anaphase-promoting complex/cyclosome (APC/C). **(k)** Endogenous co-immunoprecipitation in iPSC-derived neurons confirms BIN1 interactions with the APC/C subunits ANAPC1 and ANAPC2. Statistical significance is indicated by asterisks (\*\**p* < 0.01; \*\*\**p* < 0.001)

### 3D structural modeling of ADNeuronNet with AlphaFold3

Next, we applied AlphaFold3 (AF3) to build atomic-resolution structural models for all interactions in our ADNeuronNet (**Fig. 3d**). As controls, we extracted protein pairs within 2,075 PDB co-complex structures, and classified them into 562 interacting (positive) and 3,650 non-interacting (negative) pairs. For comparison, we further applied AF3 to 540 protein interactions from OpenCell^23^. Additionally, we also performed AF3 modeling for 4,580 randomly selected protein pairs without any prior evidence of interaction in the literature and not found within the same PDB co-complex.

We evaluated the maximum ipTM score (**Fig. 3e**) and ranking score (**Fig. S4**) among the five AF3 models generated for each protein pair. The ADNeuronNet interactions have significantly higher ipTM and ranking scores compared with non-interacting PDB pairs (Mann-Whitney U test, *p* = 1.2 × 10^−116^), random pairs (*p* = 1.7 × 10^−17^), and the OpenCell interactions (*p* = 0.021). This result further supports the high quality of our ADNeuronNet.

Furthermore, IP-MS experiments inherently cannot distinguish direct physical interactions versus co-complex associations within identified protein complexes^45^. Due to the combinatorial complexity, we would expect that most of the ADNeuronNet interactions are not direct bindings, similar to the PDB complexes that we examined above. AlphaFold3 modeling enables us not only to produce atomic-resolution structural models for ADNeuronNet interactions, but also help identify 152 interactions with high-quality models (ipTM > 0.4), which are likely direct bindings between the proteins. Among these, the interaction between WDR12 and CSNK2B has been identified through co-fractionation analysis before^46^, but it does not have any existing structure. WDR12 is a core component of the PeBoW complex, which is responsible for ribosome biogenesis, cell proliferation, and nucleolar function. CSNK2B encodes the regulatory β subunit of casein kinase II (CK2), which is a serine/threonine kinase complex that modulates many cellular signaling pathways. The AF3 structural model for this interaction has a high ipTM score (0.83) and reveals that the WDR12 residues at the predicted interface fall within WD repeats 4-6, and the CSNK2B interface residues overlap the CK2α-interaction region, both of which are known to mediate protein-protein interactions^47,48^ (**Fig. 3f**).

In fact, because of the high quality of ADNeuronNet, we were able to validate several novel interactions through endogenous co-IPs in human neurons, even when their AF3 models are of medium confidence. Here, we focused on 3 novel interactions that have never been reported previously in the literature: RIN3-SNX9 (**Fig. 3g**, ipTM= 0.40), BIN1v1-ANAPC1 (**Fig. 3i**, ipTM= 0.43), and BIN1v1-ANAPC2 (**Fig. 3j**, ipTM= 0.45). We were able to validate all 3 interactions via co-IPs using endogenous antibodies in human iPSC derived excitatory neurons (**Fig. 3h,k**). In addition, we also verified 5 other interactions reported in the literature (most likely in HEK293T cells) via endogenous co-IPs in WTC11 derived neurons and AD patient iPSC derived neurons (**Fig. S5**).

Taken together, the high quality of ADNeuronNet and the comprehensive atomic-resolution structural modeling of all interactions by AF3 provide a unique and valuable resource for elucidating underlying mechanisms of AD.

### RIN2, a neuron-specific interactor recruits BIN1 to early endosomes

BIN1, a significant AD risk gene^5^, is known to play roles in membrane remodeling, endocytosis regulation, and vesicle trafficking^49–51^. In primary neurons, BIN1 was reported to influence the endocytosis of amyloid precursor protein (APP), which can be cleaved by β-secretase in the early endosome^52,53^. Alongside its contribution to increased extraneuronal levels of Aβ, BIN1 has also been implicated in increased levels of Tau protein via its involvement in Tau spreading with extracellular vesicles^5,54,55^.

In ADNeuronNet, we identified 39 interactors (**Fig. 4a**) for the longest, neuronal isoform 1, BIN1v1, including several of its known interactors, such as AMPH and RIN3, as well as many novel interactions only observed in neurons (**Fig 4a, b**). Interestingly, while BIN1v1 has been previously reported to interact with AP2A1 and AP2A2^53,56^, in ADNeuronNet, we detected all 5 subunits of the AP-2 adaptor complex and a total of 17 proteins involved in vesicle-mediated transport^57^, most of which have never been shown to interact with BIN1 in the literature before. These results highlight the high sensitivity and quality of the ADNeuronNet pipeline.

**Figure 4.**
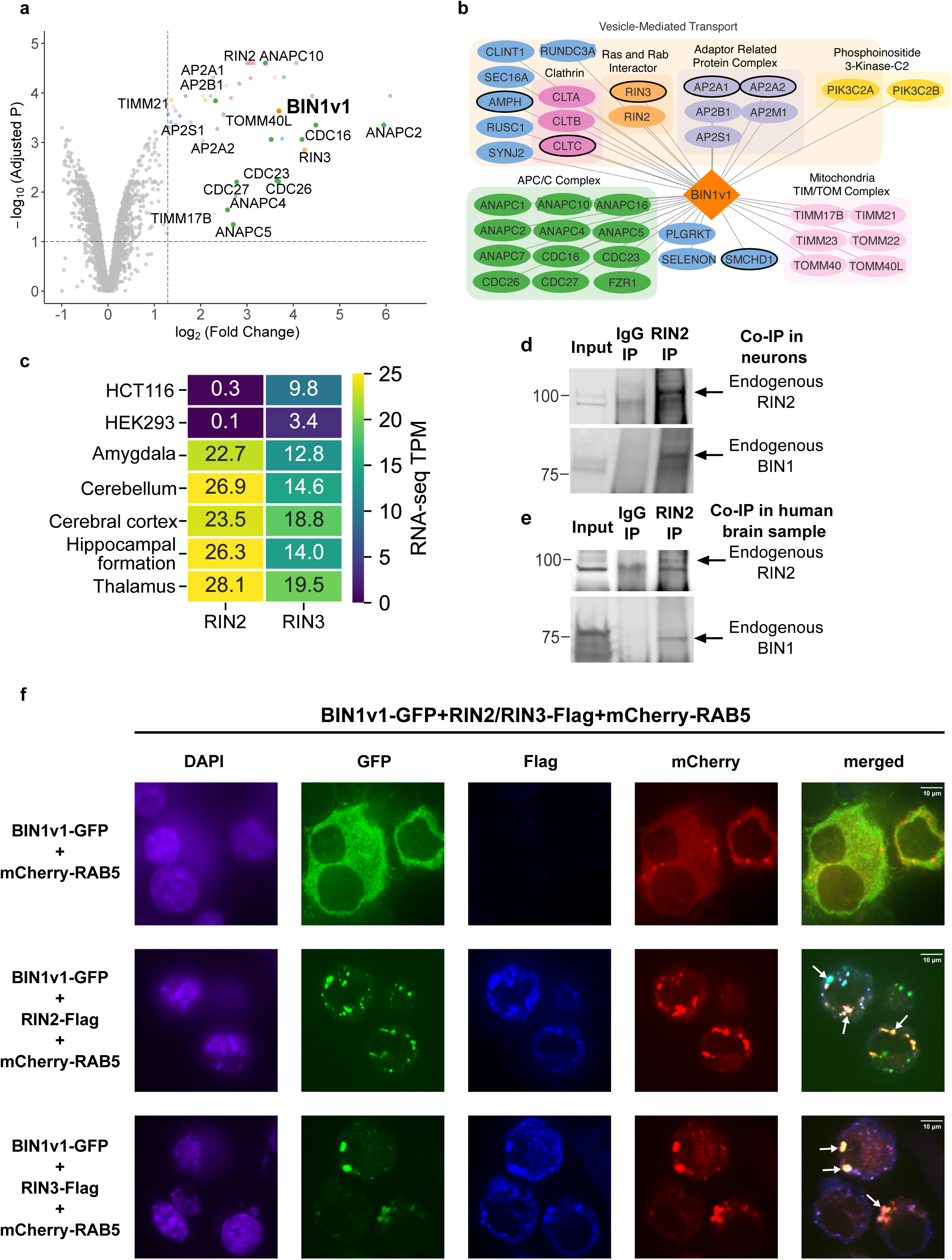
Identification and validation of a novel neuron-specific BIN1 interaction partner, RIN2. **(a)** Volcano plot of the BIN1 IP–MS experiment performed in iPSC-derived neurons. Fold changes and adjusted *p* was calculated using the analysis pipeline described in the **Methods**. Proteins with log₂(fold change) > 1.3 and adjusted *P* < 0.1 were defined as high-confidence BIN1 interactors and are highlighted in the upper right quadrant. **(b)** Network representation of BIN1 interactors identified in **(a)**, with proteins grouped according to functional similarity or known protein complex associations. **(c)** Expression levels of RIN2 and RIN3 across representative human cell lines (HCT116, HEK293) and selected human brain regions. Expression values are presented as transcripts per million (TPM). Color scale indicates absolute TPM abundance. **(d-e)** Endogenous co-immunoprecipitation western blot validation of the BIN1–RIN2 interaction in **(d)** iPSC-derived neurons and **(e)** human brain lysates. **(f)** Confocal microscopy of Neuro2a cells co-expressing BIN1v1-GFP, mCherry-RAB5 (early endosome marker), and Flag-tagged RIN2 or RIN3. DAPI stains nuclei. Arrows indicate co-localized BIN1 and RIN2/3 puncta on RAB5+ early endosomes.

One of the novel BIN1 interactors we identified is RIN2, a RAB5 guanine nucleotide exchange factor. While the interaction between BIN1 and RIN3, another Ras and Rab interactor protein, has been well established in the literature^23,53,58,59^, BIN1-RIN2 interaction has never been reported. According to Human Protein Atlas^28^, RIN2 is not expressed in HEK293T or HCT116 (thus could not have been detected in previous interactome mapping efforts for BIN1 in these two cell lines), but is highly expressed across many key brain regions (**Fig. 4c**), indicating that RIN2 may have significant roles in brain, and specifically neuronal, functions. We first validated BIN1-RIN2 interaction in iPSC derived neurons and in human brain lysate using endogenous antibodies for both BIN1 and RIN2 (**Fig. 4d, e**). Recent studies have shown that BIN1 is recruited to early endosomes through RIN3^53^, and involved in endocytic trafficking to promote APP cleavage and accumulation of Aβ peptides in the brain^53,59^. Our confocal microscopy revealed that BIN1v1-GFP is primarily cytoplasmic when expressed alone. Strikingly, co-expression with Flag-RIN2 re-localizes BIN1v1-GFP to RAB5 positive early endosomes, forming distinct puncta (**Fig. 4f**). This pattern mimics the translocation of BIN1 observed when coexpressed with RIN3^53^, suggesting that RIN2 plays a similar role to RIN3 in early endosomal AD pathology. These results highlight the power of our cell-type-specific interactome mapping in human neurons to identify key neuron-specific protein interactions that can uncover additional AD risk genes and help shed light on AD etiology.

### Patient-derived mutations alter key protein interactions associated with AD pathology

Next, we applied our highly quantitative DIA-LFQ IP-MS pipeline to examine the effects of AD-associated missense mutations on the ADNeuronNet interactions. To robustly identify changes in interaction strength among proteins, we implemented a nested interactome perturbation quantification workflow: both wildtype (WT) and mutant IPs, together with the control IPs, are all analyzed by MS in the same batch. First, we compare WT and mutant IPs to the control IPs, respectively, to identify potential interactors with the bait. Then, only for the union of the identified interactors, we compare their abundance between WT and mutant IPs to quantify changes in their interaction strength to identify strengthened (or gained) versus weakened (or lost) interactions by the mutation (**Fig. S6a**).

We collected a set of 11 high-confidence AD-associated missense mutations and isoforms (**Fig. 5a**), including well-known AD mutations from published studies^49,60–62^, prioritized GWAS variants from fine-mapping^6^, and mutations significantly associated with AD and related dementia (ADRD) from the latest Exome-Wide Association Study (ExWAS)^63^. Our list includes two BIN1 isoforms (isoform 1 and isoform 9; **Fig. 5b**) and 4 APOE alleles^64–67^ that differ at 3 amino acid sites: APOE3 (common allele; neutral; Cys112, Arg136, Arg158), APOE2 (protective; Cys112, Arg136, Cys158), APOE3CC (protective; Cys112, Ser136, Arg158), and APOE4 (most important genetic risk factor for AD^68^; Arg112, Arg136, Arg158). Among the 11 variants we examined, we discovered 19 strengthened, 9 gained, 36 weakened, and 35 lost interactions due to the mutations (**Fig. 5a**).

**Figure 5.**
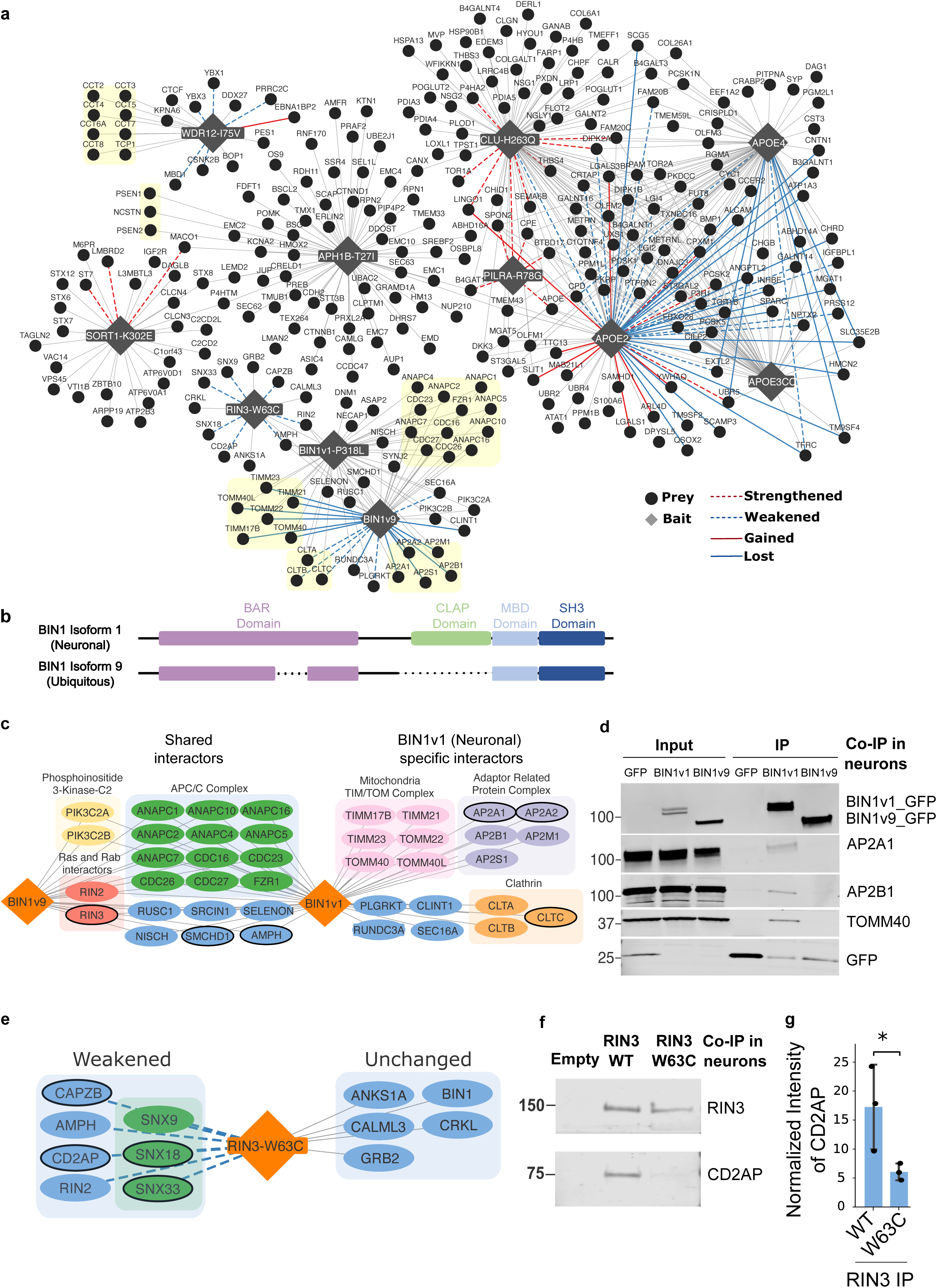
Effects of AD-associated mutations and isoforms on protein-protein interactions. **(a)** Network overview of mutation perturbation analyses. AD mutant baits are represented by grey diamonds and interactors are represented by dots. Red lines show strengthened (dashed) or gained (solid) interactions while blue lines show weakened (dashed) or lost (solid) interactions compared to the wild-type (WT). Interactions of APOE2, APOE3CC, and APOE4 alleles are compared to those of APOE3 (here considered as WT). Interactions of BIN1v9 are compared to those of BIN1v1. **(b)** Linear protein domain graphs of BIN1 isoforms 1 and 9. The dotted line indicates the isoform 1 amino acids that are missing in isoform 9. The BIN1 P318L mutation is labeled on isoform 1. **(c)** Network schematic showing shared and unique interactors of BIN1v1 and BIN1v9. The two BIN1 isoform baits are indicated by the orange diamond shapes, with shared interactors listed in between them and interactors unique to each isoform on either side of the respective bait. Proteins are colored based on functional associations or involvement in shared complexes. **(d)** Co-immunoprecipitation with anti-GFP beads followed by immunoblotting in human iPSC-derived neurons demonstrates that BIN1v1 (neuronal isoform), but not BIN1v9 (ubiquitous isoform), selectively associates with the known interactor AP2A1 and the newly identified interactors AP2B1 and TOMM40, supporting isoform-specific interactions. ‘GFP’ lanes represent neurons expressing GFP alone, and ‘BIN1v1’ and ‘BIN1v9’ lanes represent neurons expressing GFP-tagged BIN1v1 or BIN1v9, respectively. **(e)** Network representation of RIN3-W63C interaction changes compared to RIN3-WT. Proteins with weakened associations are shown on the left, whereas unchanged interactors are shown on the right. **(f)** Co-immunoprecipitation with anti-GFP beads in human iPSC-derived neurons shows decreased CD2AP binding to RIN3-W63C compared with RIN3-WT. ‘Empty’ lane represent the IP in uninfected neurons, while ‘RIN3 WT’ and ‘RIN3 W63C’ lanes represent IPs in neurons expressing GFP-tagged RIN3-WT or RIN3-W63C, respectively. **(g)** Immunoblot quantification confirms reduced CD2AP binding to RIN3-W63C compared with RIN3-WT. Statistical significance is indicated by asterisks (\**p* < 0.05)

Specifically, we compared the interactomes of two main BIN1 isoforms found in the brain^53,69^: the neuronal isoform 1 (BIN1v1) discussed previously, and the smaller, ubiquitously expressed isoform 9 (BIN1v9) (**Fig. 5b**). Studies in patient brain tissues have reported modulation of protein levels for both BIN1 isoforms in AD cases, however BIN1v1 expression was significantly reduced while BIN1v9 expression was significantly increased in AD brains compared to controls^54,70^. The defining structural difference between these isoforms is the inclusion of the CLAP (clathrin and AP-2 binding) domain in BIN1v1, which is absent in BIN1v9 ^29,69^. This difference is clearly reflected in the ADNeuronNet interactome networks for these two proteins (**Fig. 5c**). While several interactors are shared, BIN1v1 uniquely interacts with the AP-2 complex and clathrin proteins mediated by the CLAP domain^29,69^, as well as many other endocytosis- and vesicle transport-associated proteins, further confirming the high quality of ADNeuronNet. Importantly, previous large-scale human interactome mapping efforts^22,23^ used only the non-neuronal BIN1 isoform, and thus were unable to detect any CLAP-dependent interactions (**Fig. S6b, c**). Furthermore, only AP2A1 and AP2A2 interactions with BIN1v1 were reported in the literature^53,56^, whereas we were able to detect all 5 AP-2 subunits in our BIN1v1 IP, all of which are specific to BIN1v1 when compared to BIN1v9. And, we were able to confirm this binding specificity via co-IP using endogenous antibodies in neurons (**Fig. 5c, d**). These results confirm the high sensitivity and specificity of our neuron interactome mapping pipeline.

Additionally, we detected subunits of the mitochondrial transport TIM/TOM complex interacting with BIN1v1 in ADNeuronNet, which have not been previously reported with any BIN1 isoform in the literature and did not appear in the BIN1v9 network (**Fig. 5c**). We were able to again confirm this novel, specific interaction between BIN1v1 and TIM/TOM using co-IP in neurons (**Fig. 5d**). Certain TIM/TOM subunits, namely TOMM40, the central channel of the TOM complex^71^, TOMM22, and TIMM23^72^, have been identified to participate in mitochondrial accumulation of Aβ peptides, which can cause toxicity in the mitochondria by entering through the TOMM40 pore, or can clog the TOM complex to inhibit proper protein import, both of which may lead to mitochondrial dysfunction in AD^71,72^. The link between mitochondrial transport and BIN1 has not yet been characterized, however our results suggest that this connection may be specific to a neuronal context.

Furthermore, an AD patient variant that caused notable perturbations in the protein interaction network was a RIN3 missense mutation rs150221413 (p.Trp63Cys) discovered in a whole exome sequencing study of 93 early-onset AD patients^62^. It is located in the SH2 domain of RIN3 and is predicted to be deleterious by multiple prediction tools (CADD = 23.60; Polyphen = 1.00; SIFT = 0; REVEL = 0.52). The mechanism of how this mutation contributes to AD risk is not yet clearly understood, however it is believed that increased expression of both WT RIN3 or the p.Trp63Cys variant can contribute to disease development^73^. As discussed above, RIN3 recruits BIN1 and CD2AP to Rab5-positive early endosomes, forming a complex that can affect APP trafficking and processing into Aβ peptides^59^. Interestingly, we found that although the interaction with BIN1 remains unchanged between the WT and p.Trp63Cys mutant, the interaction between RIN3 and CD2AP is significantly weakened by the p.Trp63Cys mutations. To confirm our findings, we performed co-IP in iPSC-derived neurons expressing GFP-tagged WT or p.Trp63Cys RIN3. The blot was probed with an anti-CD2AP antibody to detect endogenous CD2AP interacting with RIN3. After normalizing for protein expression, RIN3 p.Trp63Cys exhibited a ∼3-fold reduction in CD2AP binding compared to WT in three independent co-IP experiments (**Fig. 5f, g**). A previous study has reported that depletion of CD2AP can lead to the failure of APP to be sorted for degradation, resulting in enhanced APP and BACE1 convergence in early endosomes^52^. Additionally, we noticed that the RIN3 mutation caused weakened interactions with sorting nexin proteins, SNX9, SNX18, and SNX33. These proteins are paralogs and the only members of the sorting nexin family that contain an SH3 domain^74^. Both SNX9 and SNX33 have been found to interact with dynamin, establishing their connections to endocytic pathways.^75^ Taken together, RIN3 p.Trp63Cys mutation weakened several key interactions and could potentially contribute to AD risk through affecting multiple processes.

Altogether, our quantitative interactome perturbation studies in human iPSC-derived neurons reveal that the AD-associated isoforms and mutations can strengthen/gain or weaken/lose interactions with key proteins. These results can be utilized to direct future investigations on the specific roles that these isoforms and mutations play in neuronal functions and AD progression.

### Novel BIN1 interaction with APC/C complex reveals insights into Tau pathobiology

Proteomic analysis of the BIN1 interactome identified multiple previously unrecognized binding partners in addition to RIN2, including twelve subunits of APC/C, a multi subunit E3 ubiquitin ligase with established roles in neuronal proteostasis and synaptic maintenance^76^. We focused on two core APC/C components, ANAPC1 and ANAPC2, to validate this interaction. Co-IP experiments in human iPSC derived neurons confirmed that endogenous BIN1 associates with both ANAPC1 and ANAPC2 (**Fig. 3k**).

This interaction is particularly relevant in the context of human Tau pathology. In the postmortem human brain, BIN1 protein levels correlate with neurofibrillary tangle burden but not with diffuse or neuritic amyloid plaques^54^. Consistent with this observation, the BIN1 rs744373 risk variant is associated with increased Tau PET signal across brain regions vulnerable to Tau accumulation, without corresponding changes in amyloid PET^55^. Moreover, GWAS on 11 ADRD-related neuropathology endophenotypes identified Braak NFT stage is strongly associated with BIN1 rs6733839 risk variant^77^.

Based on the identification of APC/C components as BIN1 interactors, we next examined whether disruption of APC/C function influences Tau pathology in human neurons. ANAPC1 or ANAPC2 expression was selectively reduced using shRNA, with efficient knockdown confirmed by immunoblotting (**Fig. 6b, c**). Depletion of either subunit did not grossly alter neuronal morphology, although sustained ANAPC2 knockdown resulted in a modest but significant reduction in MAP2 immunoreactivity (**Fig. 6d, f**). Tau aggregation was induced using Tau fibrils (K18) and quantified by MC1 immunostaining. Knockdown of either ANAPC1 or ANAPC2 significantly increased seeding-induced MC1 immunoreactivity relative to scrambled control shRNA (Sh-Ctrl), indicating enhanced Tau propagation following deficiency of either BIN1 interacting APC/C subunit (**Fig. 6d, g**).

**Figure 6.**
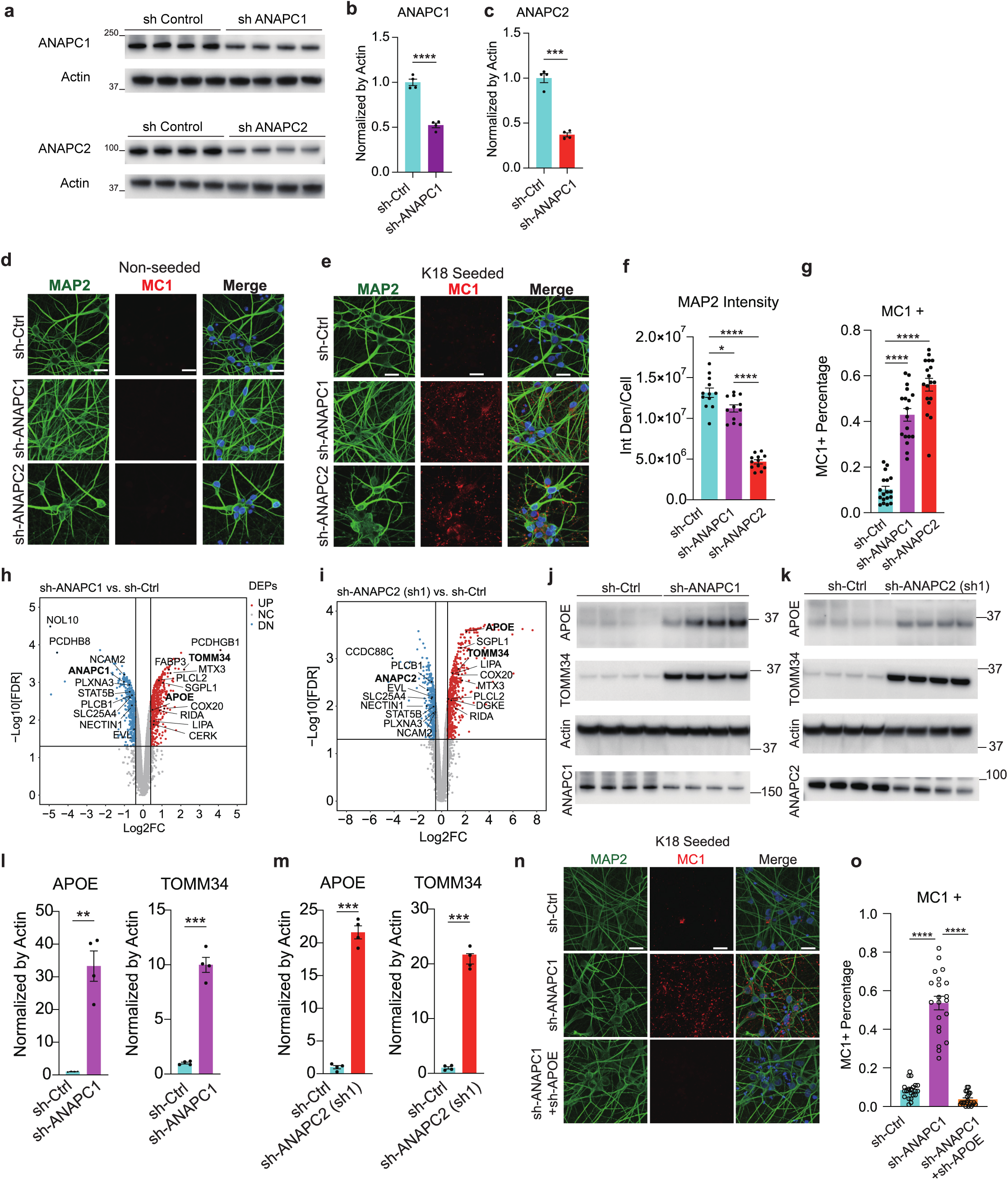
Loss of BIN1 associated APC C complex components enhances Tau aggregation by increasing neuronal APOE. **(a–c)** Immunoblot (a) and quantification confirms efficient knockdown of ANAPC1 (b) and ANAPC2 (c) following shANAPC1 or shANAPC2 transduction in human 4R P301S neurons. **(d–e)** Representative immunostaining of MAP2 (green) and the pathological Tau conformer MC1 (red) in human 4R P301S neurons under unseeded (d) and Tau seeded (e) conditions. **(f–g)** Quantification of MAP2 (f) and MC1 (g) signals in neurons transduced with shANAPC1 or shANAPC2. Knockdown of either ANAPC1 or ANAPC2 selectively increased seeding induced MC1 immunoreactivity without affecting neuronal MAP2 levels. **(h–i)** Volcano plots from quantitative proteomic analyses comparing lysates from neurons transduced with shCtrl versus shANAPC1 (h) or shANAPC2 (i). **(j–m)** Immunoblot and quantification demonstrate that shANAPC1 reduces ANAPC1 and increases APOE and TOMM34 levels (j, l), while shANAPC2 reduces ANAPC2 and similarly increases APOE and TOMM34 levels (k, m) in 4R P301S neurons. **(n–o)** Representative immunostaining and quantification of MC1 in Tau seeded neurons transduced with shANAPC1 alone or in combination with shAPOE. Co depletion of APOE abolishes the shANAPC1 induced increase in MC1 signal, indicating that elevated neuronal APOE mediates Tau aggregation downstream of APC/C disruption. Statistical significance is indicated by asterisks (\*\**p* < 0.01; \*\*\**p* < 0.001; \*\*\*\**p* < 0.0001)

To define molecular pathways linking APC/C disruption to enhanced Tau propagation, we performed unbiased quantitative proteomic analyses in human neurons following ANAPC1 or ANAPC2 knockdown. Both perturbations produced highly concordant proteomic changes, consistent with a shared downstream response to APC/C impairment (**Fig. S7b,c**). Proteins involved in neurogenesis and neuronal development were consistently downregulated following depletion of either APC/C subunit, including PLCB1, NCAM2, NECTIN1, EVL, PLXNA3, SLC25A4, and STAT5B, suggesting compromised neuronal maintenance and resilience (**Fig. 6h, i; Fig. S7a**). In contrast, APC/C disruption led to coordinated upregulation of proteins associated with mitochondrial function and lipid metabolism. Mitochondrial associated proteins such as COX20, TOMM34, and MTX3 were consistently increased, alongside a prominent induction of lipid handling pathways. Notably, APOE emerged as one of the most strongly upregulated proteins, together with FABP3, LIPA, CPT2, CERK, DGKE, and SGPL1 (**Fig. 6h, i; Fig. S7a**).

Venn diagrams confirmed robust overlap among upregulated proteins across ANAPC1 and ANAPC2 knockdown conditions (**Fig. S7b**). Similarly, substantial overlap was observed among downregulated proteins induced by shANAPC1 and two independent shANAPC2 constructs (**Fig. S7c**). Pathway enrichment analysis of the shared proteomic changes further supported these findings, revealing significant enrichment of lipid metabolism and mitochondrial pathways among upregulated proteins, and neuronal development and synaptic related pathways among downregulated proteins (**Fig. S7d, e**).

To validate the proteomic results and establish a link between APC/C disruption and APOE induction, we performed immunoblot analyses in human neurons transduced with shANAPC1 alone or in combination with shRNA targeting APOE. These experiments confirmed efficient knockdown of ANAPC1 and demonstrated corresponding modulation of APOE protein levels, validating APOE as a downstream target of APC/C disruption (**Fig. S7f, h**).

The robust and reproducible induction of APOE prompted us to test whether neuronal APOE mediates Tau aggregation downstream of APC/C dysfunction. Immunoblot analysis confirmed increased APOE and TOMM34 protein levels following ANAPC1 or ANAPC2 knockdown (**Fig. 6j–m**). Importantly, lentiviral mediated depletion of APOE markedly attenuated the increase in MC1+ Tau inclusions induced by ANAPC1 loss, effectively rescuing the Tau propagation phenotype (**Fig. 6n, o**). Together, these data identify a previously unrecognized BIN1–APC/C–APOE regulatory axis in human neurons and demonstrate that loss of APC/C function promotes Tau aggregation through aberrant induction of neuronal APOE. This mechanism provides a direct molecular link between BIN1 associated proteostasis pathways and Tau pathology associated with BIN1 protein levels and genetic risk^54,55,77^.

## Discussion

Current AD-specific protein interaction mapping is generally limited in scope, while large-scale interactomes were not performed in AD-relevant brain cell types. In this study, we mapped interactions for 61 AD risk proteins in iPSC-derived human neurons to create the first large-scale neuron-specific AD protein interactome, ADNeuronNet, consisting of 1,769 interactions among 1,145 proteins. The high quality of ADNeuronNet is reflected in its significant overlap of known interactions from the literature. Furthermore, we observed significantly higher percentages of neuron-expressed and neuron-specific interactors in ADNeuronNet compared to known interactions in the literature, confirming the neuron-specific nature of our interactome. Additionally, the high overlap of ADNeuronNet interactors with Tau-inclusion-associated CRISPRi hits signified its relevance in a disease context. Finally, we generated atomic-resolution structural models for all ADNeuronNet interactions using AlphaFold3, and found the ADNeuronNet interactions had significantly higher ipTM scores compared to controls.

Although TREM2 and APOE are predominantly expressed in microglia and astrocytes, their disease-relevant effects converge on excitatory neurons, the principal sites of synaptic failure and degeneration in AD. Both pathways generate extracellular or internalized signals that directly engage neuronal protein networks, motivating interrogation of their interactomes within neurons rather than inference from expression alone. Proteolytic shedding of TREM2 produces soluble TREM2^78^, a bioactive ligand that exerts neuroprotective functions and can be taken up by neurons^79^. These TREM2-derived species are positioned to interact with neuronal proteins governing endosomal trafficking, lysosomal function, and synaptic homeostasis. Similarly, astroglia-derived APOE is avidly internalized by neurons, where it traffics through endolysosomal compartments central to Tau propagation and synaptic integrity^80^. In addition, prior studies established a critical role of neuronal APOE modulates neurotoxicity in a cell autonomous manner^81^. In this study, we further identified the BIN1-ANAPC1-APOE axis in Tau propagation in excitatory neurons.

We demonstrated how ADNeuronNet can be utilized to identify novel, neuron-specific proteins with potential to impact AD pathology. In particular, we focused on BIN1, the most predominant AD risk gene that is associated with Tau, not amyloid pathology; yet, the mechanisms are still unknown^82^. In ADNeuronNet, we found RIN2, a protein not expressed in HEK293T or HCT116 cell lines, as a novel interactor of BIN1. When co-expressed with BIN1, RIN2 recruited BIN1 to early endosomes in a similar fashion to known BIN1 interactor and AD risk protein, RIN3. This suggests a similar mechanism between RIN2 and RIN3 underlying AD pathogenesis. Interestingly, RIN2 is highly expressed (even higher than RIN3) in the cerebral cortex, hippocampus, and other key brain regions. We also identified novel BIN1 interactions with APC/C complex proteins. Strikingly, we found that loss of ANAPC1 or ANAPC2 via an shRNA knockdown resulted in increased levels of Tau inclusions in neurons. Proteomic profiling of these cells revealed APOE was upregulated in both ANAPC1 and ANAPC2 knockdown conditions compared to the control. When the ANAPC1 knockdown was followed by APOE knockdown in the same cell line, we observed dramatic reduction of Tau aggregation, producing a near-complete rescue. These findings identify a previously unrecognized BIN1-APC/C-APOE regulatory axis as a novel mechanism by which BIN1 promotes Tau propagation in human neurons, linking BIN1 and APOE, two most prominent AD risk genes.

Finally, we showed that our pipeline can be used to quantitatively detect changes in protein interactome networks caused by AD-associated isoforms or mutations. For example, we found that the AP-2 complex and clathrin proteins specifically interact with the neuronal isoform 1 of BIN1, but not the smaller, ubiquitous isoform 9, in agreement with previous studies^29,69^. Furthermore, in ADNeuronNet, we also detected and validated with endogenous co-IP in neurons that the mitochondrial TIM/TOM complex interacts with BIN1v1 (which has never been reported in the literature before), but not BIN1v9. By comparing the interaction networks of WT and disease-associated mutations, we are able to gain further insight into how these mutations contribute to AD progression. Of note, we successfully carried out comparative interactome mapping for RIN3 W63C mutation and identified specific interaction rewiring by this AD mutation. We validated the disruption of its binding to another key AD protein, CD2AP via co-IP in human neurons.

Although iPSC-derived neurons certainly have limitations to recapitulate aging and the complexity of cell-cell interactions in human brains, using homogenous human neurons for our IP-MS experiments also offers several unique advantages. First, human brain samples always contain a mixture of many different cell types. To date, it is impossible to obtain brain samples of a specific cell type for IP-MS experiments at scale. Therefore, it is simply impossible to generate large-scale cell-type-specific interactome maps using human brain samples. Furthermore, the composition of the cell types also varies drastically from sample to sample, making it extremely difficult, if not impossible, to separate changes in protein interactions due to sample composition differences versus underlying disease risks. Not to mention, significant gene regulation changes occurring post-mortem in brain tissues^83^ may additionally impact protein interaction results.

Furthermore, our extensive *in vivo* validation and functional studies with endogenous proteins confirm that our ADNeuronNet indeed captures functionally relevant protein interactions within human neurons. Endogenously tagging each bait through CRISPR knock in^84–86^ is still quite time consuming, and does not yet scalable to carry out large-scale interactome mapping efforts for human neurons or other brain cell types. However, for the 40 neuron-expressed bait genes that failed cloning due to their large size, endogenous tagging provides a viable alternative that we are actively pursuing to complete our neuron interactome map.

Beyond throughput, our lentiviral-infection based workflow actually has unique advantages to study interactome differences between isoforms and perturbations caused by disease-associated mutations in a high-throughput fashion, which are either highly difficult (for isoforms) or extremely time consuming (through base/prime editing for mutations) for endogenous tagging based approaches. As we have shown in our ADNeuronNet, different isoforms of the same protein can have quite distinct interactomes and thus have varying functions. Furthermore, many mutations in key AD genes can significantly impact disease risk. For example, APOE3, the most common allele of APOE in the population, has no influence on AD risk, while APOE4 is the most significant genetic AD risk factor, and APOE2 and 3CC are regarded to have protective abilities^64–67^. directly comparing protein interactors of mutated baits to their wild-type can provide illuminating information on the specific biological pathways the AD-associated mutation impacts. This can help to elucidate the molecular mechanisms that link the mutation to AD pathogenesis. Altogether, the key advantages to our pipeline make it an effective and optimized system for AD protein interactome mapping.

In conclusion, the broad scope and high level of depth offered by ADNeuronNet make it a valuable resource for guiding future AD research and for studying how proteins function in neurons and in human brains in general. Subsequent investigations into the pathological roles of ADNeuronNet interactions can lead to the discovery of vital disease mechanisms and potential therapeutic targets. This is especially important given that a massive amount of DNA sequencing data have been accumulated across traits, including for AD. However, traditional GWAS and statistical methods are often underpowered and require a very large sample size. Furthermore, statistical analysis of variants does not identify a specific change in protein function; thus it may not be sufficient to accurately define causal variants, and often ignore those that are very rare. Therefore, it is paramount to integrate such statistical analyses within the protein-protein interactome framework to borrow strength from neighboring genes in the network and to infer functional impact of variants. Our ADNeuronNet, being the first interactome map in human excitatory neurons, will be a critical resource for achieving these goals not only for AD research, but for many other neurological and neuropsychiatric disorders in general.

Future studies should expand our neuron-specific interactome mapping to encompass additional genes implicated in AD risk, as revealed by our ADNeuronNet – particularly hub interactors that connect with multiple AD risk proteins and may represent points of pathway convergence. Although we examined seven AD-associated mutations as a proof of principle to demonstrate the specificity and sensitivity of our platform, a more comprehensive, systematic interactome perturbation study of a broader spectrum of disease-linked variants will be essential to uncover shared molecular mechanisms and converging pathogenic pathways. Finally, our ADNeuronNet paves the way for large-scale interactome mapping in other key brain cell types, especially microglia, which express many AD risk genes and are strongly implicated in AD pathogenesis^87^. Further extending this approach to multicellular human cerebral organoid systems^88,89^ may further enable interrogation of protein interactions within a more physiologically relevant context.

## Methods

### Selection and Cloning AD Risk Genes

As described above, we included all 86 high-confidence protective or causal genes with genetic and functional evidence compiled by Alzheimer’s Disease Sequencing Project (ADSP), as well as all 114 genes from the 3 latest AD GWAS studies^5,26,27^. These studies prioritized 89 risk genes from GWAS loci through comprehensive and advanced analyses, including position mapping, colocalization of molecular QTL data (expression, splicing, protein levels, methylation, and/or histone acetylation) from AD-relevant tissues, statistical fine-mapping, drug target information, and previous literature. For comprehensiveness, all nearest genes to all significant lead SNPs were also included (25 additional genes)^5,26,27^. In total, we compiled a list of 137 unique AD-associated genes with strong genetic support.

Among the 137 unique AD-associated genes, 97 were classified as neuron-expressed based on integrated analysis of ENCODE transcriptomic datasets and our in-house RNA-seq data from iPSC-derived neurons. Specifically, transcript abundance (TPM) was quantified for pyramidal, bipolar, and motor neurons using 13 publicly available ENCODE RNA-seq datasets, together with RNA-seq data from iPSC-derived neurons from our previous study^31^. For each gene, TPM values were averaged across these neuronal datasets. Genes with an average TPM greater than 1 were defined as neuron-expressed. To define neuron-specific genes, we applied a more stringent criterion: mean neuronal TPM > 5 across all neuronal datasets, combined with normalized TPM < 1 in both HCT116 and HEK293 cells, as determined using expression data from the Human Protein Atlas.

Each neuron-expressed AD risk gene sequence was initially cloned into a FUGW (Addgene, 14883) vector, resulting in a C-terminal GFP tagged protein bait for downstream IP-MS analysis; genes that failed the initial IP-MS experiment, were re-cloned into the FUGW plasmid with an N-terminal GFP tag.

### Human iPSC Culture and Neuron Differentiation

The i^3^N iPSCs were cultured without a feeder layer on Matrigel-coated (Corning, 354277) plates in mTeSR Plus medium (StemCell, 100-0276). The cells were passaged with Accutase (StemCell, 7920) every 3-4 days, and initially seeded with ROCK inhibitor Y-27632 (StemCell, 72304) on the first passage followed by a media change the next day to remove the Y-27632.

The i^3^N iPSCs were differentiated into i^3^Neurons using the two-step protocol as described in Wang et al^30^. During the pre-differentiation step, the i^3^N iPSCs were incubated with doxycycline (2 μg/mL, Millipore Sigma, D9891) in knockout DMEM (KO-DMEM)/F12 medium (Gibco, 12660-012) with 1X N-2 supplement (Thermo Fisher Scientific, A3890401), 1x non-essential amino acids (NEAA), mouse laminin (0.1 μg/mL, Thermo Fisher Scientific, 23017015), brain-derived neurotrophic factor (BDNF, 10 ng/mL, PeproTech, 450-02), neurotrophin-3 (NT3, 10 ng/mL, PeproTech, 450-03), and Y-27632 for 3 days with the medium changed daily and Y-27632 removed from day 2. During the maturation step, the pre-differentiated cells were dissociated with Accutase and subplated on poly-D-lysine (PDL)-coated plates in 50% DMEM/F12 medium (Gibco, 11330032), 50% Neurobasal Plus medium (Gibco, A3582901), 0.5x B27 supplement (Thermo Fisher Scientific, A3582801), 0.5x N-2 supplement, 1x NEAA, 1x GlutaMAX (Thermo Fisher Scientific, 35050061), mouse laminin (0.1 μg/mL), BDNF (10ng/mL), NT3 (10 ng/mL), and doxycycline (2 μg/mL). Every 3 days, half the medium was replaced. Once the differentiation period was complete, the cells were harvested by washing once with cold PBS, then pelleting for downstream IP-MS experiments.

### Lentiviral Transduction

Lentivirus was produced in human embryonic kidney 293T (HEK293T) cells cultured in Dulbecco’s modified Eagle’s medium (DMEM) (Gibco, 11965-118) supplemented with 10% FBS (Avantor, 97068-085). The 293T cells were seeded at a density of 0.4 × 10^6^ cells per well in 2 wells of a 6-well plate and incubated at 37°C overnight. The next day, the medium was replaced with 3 mL of fresh medium per well immediately before the cells were transfected via a polyethylenimine (PEI) (1 mg/mL, Polysciences, 23966) transfection containing a mixture of the following plasmids in 200 μL of Opti-MEM (Gibco, 31985-062) per well: 1.14 μg FUGW plasmid containing the gene of interest, 0.57 μg psPAX2 (Addgene, 12260), and 0.285 μg pMD2.G (Addgene, 12259). After 72 h from transfection, then the supernatant containing the virus was collected from both wells, combined, and filtered through a 0.45 µm PES filter. The virus was concentrated using Lenti-X Concentrator (Takara, 631231) overnight at 4°C. The next day, the virus was centrifuged at 3000 × g for 1 h at 4°C to obtain the viral pellet. The supernatant was discarded and the viral pellet was resuspended in 100 μL of cold, filtered PBS, then used immediately for infection or frozen in −80°C for storage.

One day before lentiviral transduction, i^3^N iPSCs were seeded with mTEsR Plus medium and Y-27632 at a density of 0.1 × 10^6^ cells in a matrigel-coated well in a 12-well plate. One hour before transduction, the medium was replaced without Y-27632. The virus resuspended in PBS was added dropwise to the i^3^N iPSCs and incubated at 37°C overnight. The following day, the virus-containing media was replaced with fresh medium without Y-27632 and the cells were proliferated for sorting.

### FACS

For fluorescence-activated cell sorting, GFP-expressing i^3^N iPSCs were dissociated with Accutase, pelleted, resuspended in PBS containing DAPI (4’,6-Diamidino-2-Phenylindole, Dihydrochloride, 1 mg/mL, Invitrogen, D1306) diluted 1:1000, and filtered. The isolated cell suspensions were sorted with a BD FACSMelody Cell Sorter. Gates for scatter (FSC-A vs SSC-A) and single cells (FSC-H vs FSC-A) were applied. Additional gates for DAPI and GFP were used to isolate live cells with high GFP-expression. Sorted cells were collected in a matrigel-coated 24-well plate containing mTeSR Plus medium with Y-27632 and proliferated until ready for neuron differentiation.

### Immunoprecipitation of GFP-tagged Proteins

The cell pellets were resuspended in 500 μL of Digitonin lysis buffer (50 mM HEPES pH 7.5, 150 mM KCl, 2 mM MgCl^2^, 1.5% Digitonin (high purity, Calbiochem), 0.1% Pierce Universal Nuclease (Thermo Fisher Scientific, 88700)) with freshly added cOmplete protease inhibitor cocktail (Roche, 11873580001), PMSF (Sigma-Aldrich, 93482), and PhosSTOP (Roche, 4906837001), then incubated at 4°C for 1 h with rotation. The whole cell lysates were sonicated in a cup horn sonicator (Branson) for 30 sec at 40% amplitude, then centrifuged at 21,000 × g for 15 min at 4°C. The supernatants containing the cleared extracts were collected and split into 4 separate replicate IPs. GFP Selector magnetic beads (NanoTag Biotechnologies, N0310) (5 μL slurry per IP) were washed 3 times with wash buffer (50 mM HEPES pH7.5, 150 mM KCl, 2 mM MgCl^2^) and added to the cleared lysate, then rotated overnight at 4°C. The protein-bound beads were washed twice with 800 μL of wash buffer, then resuspended in 100 μL of elution buffer (100 mM HEPES pH 7.5, 1% SDS) and incubated at 95°C for 7 min. Using a magnetic stand, the eluates were separated from the beads and collected in a new tube.

### Proteomic Sample Preparation

Whole proteome samples were produced by lysing cell pellets with cold RIPA buffer (50 mM Tris pH7.4, 150 mM NaCl, 1% NP-40, 0.1% SDS, 0.5% sodium deoxycholate) supplemented with protease inhibitor (Roche, 11873580001), PMSF (Sigma-Aldrich, 93482), and PhosSTOP (Roche, 4906837001). The lysates were rotated at 4°C for 30 min, sonicated in a cup horn sonicator (Branson) for 90 sec at 40% amplitude, then cleared via centrifugation at 21,000 × g for 15 min at 4°C. Supernatants were transferred to a new tube and protein concentrations were measured with Bio-Rad Protein Assay (Bio-Rad).

Both whole proteome samples and IP eluates were reduced using 200 mM TCEP (Thermo Fisher Scientific, 77720) incubated at room temperature for 30 min, then alkylated with 375 mM iodoacetamide incubated at room temperature for 30 min, protected from light. Sample proteins were precipitated and washed 3 times with a precipitation solution (50% acetone, 49.9% ethanol, 0.1% acetic acid). The protein pellets were resuspended in 8 M Urea, which was then diluted to 2 M Urea by adding NaCl/Tris solution (50 mM Tris pH 8.0, 150 mM NaCl). The samples were digested with 500 ng of mass-spectrometry grade Trypsin Gold (Promega, V5280) and nutated overnight at 37°C. The digestion was quenched by adding equal volume of 4% formic acid. The peptide samples were desalted by either loading onto Evotips according to the manufacturer’s protocol (Evosep, EV2011), or with lab-made C18 solid-phase extraction spin columns. Briefly, C18 sorbent from a Sep-Pak 200 mg C18 (Waters, WAT054945) was distributed into empty micro-spin columns (Thermo, 89879), which were conditioned and equilibrated with 80% acetonitrile in 0.1% acetic acid and 0.1% TFA, respectively by spinning at 1000 × g for 1 min each step. The samples were added to the spin columns, washed with 0.1% acetic acid, eluted with 80% acetonitrile in 0.1% acetic acid and dried in a Speed-Vac. Immediately before mass spectrometry analysis, the dried samples were resuspended in 0.1% TFA.

### Liquid Chromatography- Mass Spectrometry Acquisition

Samples were analyzed with either a nanoElute2 (Bruker) or Evosep One (Evosep) liquid chromatography system coupled to a timsTOF HT mass spectrometer (Bruker). For the nanoElute2, a two-column separation method was employed, including a 5 mm PepMap Neo trap Cartridge (Thermo Fisher Scientific, 174500) and an in-house 10 cm by 100 µm analytical column packed with 1.9 µm C18 beads (Dr. Maisch, r119.aq.0001). Peptide separation for the nanoElute2 was performed over a 25 minute elution gradient consisting of solvent A (0.1% formic acid in water) and steadily increasing concentration of solvent B (0.1% formic acid in acetonitrile) from 2% to 35% B. For the Evosep One, an in-house 8 cm by 150 µm capillary column packed with 1.9 µm C18 beads was used with no trap column. The standard Evosep 60 SPD method was used for peptide separation. Sample acquisition was performed in data-independent acquisition mode with parallel accumulation-serial fragmentation (dia-PASEF), employing 12 cycles with estimated cycle time of 1.7 seconds. In total, 34 mass windows of 25 Da width and 2 mobility windows each were acquired with a ramp time of 75 ms to cover a mass range from 350 to 1250 Da and a mobility range from 0.64 1/K0 [Vs/cm-2] to 1.37 1/K0 [Vs/cm-2]. Collision energy was set as a function of ion mobility to follow a straight line from 20 eV at 0.6 1/K0 [Vs/cm-2] to 59 eV at 1.6 1/K0 [Vs/cm-2]. Prior to running the samples, the elution voltage calibration was confirmed, and if needed, calibrated for 1/K0 ratios using three ions from ESI-L Tuning Mix (Agilent, G1969-85000) (m/z 622, 922, 1222) in the timsControl software (Bruker).

### Spectral Library Generation

Lyophilized peptides from 250 μg protein of i^3^Neuron lysates were resuspended in 15 μL LC-grade H2O, 10 μL 10% FA, and 60 μL HPLC-grade acetonitrile. A total of 80 μL was injected into an UltiMate 3000 system and fractionated by HILIC (hydrophilic interaction liquid chromatography) using a TSKgel Amide-80 column (2mm×150mm, 5 μm; Tosoh Bioscience). The following gradient buffers were used: buffer A (90% acetonitrile), buffer B (75%acetonitrile and 0.005% TFA), and buffer C (0.025% TFA). The gradient program was as follows: 100% buffer A (0 min); 98% buffer B/2% buffer C (5 min); and 5% buffer B/95% buffer C(35 – 40 min). Flow rate: 150μL/min. Fractions were collected every 30 seconds from minute 5 to minute 23. Fractions 27 to 36 were pooled together. All fractions were dried, resuspended in 10 μL of 0.1P, split into two replicates for LC-MS/MS analysis and subjected to both DDA and DIA acquisition. DIA runs and DDA runs of fractions were searched using Fragpipe v22.0 to build the spectral library.

### IP-MS Data Analysis

Raw mass spectrometry data acquired in data-independent acquisition (DIA) mode were processed using DIA-NN (version 1.8.1). A reviewed UniProt human protein sequence database (Swiss-Prot, November 2023 release; taxonomy ID 9606) was used for all searches. Spectral libraries generated as described above were used where applicable. In experiments in which the bait protein was not detected using the specific spectral library—owing to insufficient or absent representation of bait-derived peptides—DIA-NN’s library-free search mode was employed. This strategy enabled unbiased peptide identification directly from DIA data and was essential for robust bait detection and quantification in the absence of an appropriate project-specific spectral library. For DIA-NN searches, up to three missed cleavages and a maximum of two variable modifications per peptide were allowed. Carbamidomethylation of cysteine residues and oxidation of methionine were specified as fixed modifications. The fragment ion m/z range was set from 100 to 1800. Match-between-runs (MBR) was enabled, and peptide quantification was performed using the “Any LC (high accuracy)” strategy.

DIA-NN output data were further analyzed and visualized using Python and R. Protein enrichment was assessed relative to the GFP control using four biological replicates. For each protein, abundance ratios were computed by comparing IPs vs GFP controls, and fold change (FC) was calculated as the median of all possible ratios associated with that protein. Statistical significance (i.e., raw *p*) was evaluated using an empirical Bayes framework implemented in the limma package, which is particularly well suited for proteomics datasets with limited replicate numbers. This approach stabilizes variance estimates by borrowing information across all quantified proteins, thereby improving statistical power and robustness while reducing false positives arising from noisy measurements. Adjusted *p* was generated using the Benjamini–Hochberg procedure to adjust for multiple hypothesis testing and control the false discovery rate (FDR). Proteins commonly identified as contaminants across many baits, including keratins, ribosomal subunits, SURF6, VGF, and SYP, were excluded from downstream analyses. For a protein to be considered an interactor of the bait, we require log2FC > 1, adjusted *p* < 0.1, and precursor count ≥ 8. We manually inspected each volcano plot for all of our IPs; for some baits, we applied even more stringent log2FC and adjusted *p* cutoffs optimized for that particular bait IP for the final high-confidence interactome.

To quantify differences in protein interactions between wild-type and mutant conditions, we developed a nested analytical workflow (**Fig. S6**). Protein interactors for each condition were first identified by comparing wild-type or mutant bait IPs to corresponding GFP controls, applying thresholds of FC > 1.5 and FDR < 0.1. To minimize noise, subsequent analyses only focused on the union of interactors detected in either the wild-type or mutant bait IPs. To evaluate mutation-specific effects on protein–protein interactions, a direct comparison was performed between mutant and wild-type bait IPs. Protein intensities were normalized to the abundance of the bait protein in each sample to account for differences in bait expression levels across conditions. Results were visualized using volcano plots, in which log2FC were plotted against adjusted *p*. Proteins exhibiting differential interactions between mutant and wild-type conditions were defined as |log2FC| > 0.58 (i.e., FC > 1.5), adjusted *p* < 0.1 (**Fig. S6**).

### Enrichment analysis of ADNeuronNet

To evaluate the quality of ADNeuronNet, we assessed enrichment relative to published human interactomes, restricting analyses to interactions involving shared baits with our ADNeuronNet. Published human interactome datasets included: (i) Literature-PPI, comprising all known protein-protein interactions from the literature: In our HINT^90^ database, we compiled a comprehensive set of experimentally validated protein-protein interactions for human (all-quality, combined binary and co-complex interactions), by integrating information from commonly used databases, including BioGRID^91^, DIP^92^, IntAct^93^, MINT^94^, iRefWeb^95^, HPRD^96^, PDB^97^, and MIPS^98^. We also made sure to include all published interactions from BioPlex^22,99,100^ and OpenCell^23^; (ii) BioPlex; (iii) OpenCell. Enrichment was calculated using Fisher’s exact test, reported as log odds ratios (logOR), with standard errors estimated using the delta method.

To assess functional relevance to Tau pathology, we examined overlap between ADNeuronNet interactors and Tau inclusion-associated CRISPRi hits identified in 4R-P301S-dCas9 iPSC-derived neurons. Interactors were stratified by the number of AD risk baits they bound (all; ≥2; ≥3; ≥4), and overlap proportions were compared to literature-based interactors using a two-proportion Z-test.

Disease-relevance enrichment was evaluated using differentially expressed genes (DEGs) from a new ROSMAP snRNA-seq dataset (∼1.3 million nuclei; 48 individuals; six brain regions)^35^. DEG calls were defined by the original study, requiring concordant significance and direction in both Nebula and MAST. Enrichment of ADNeuronNet interactors in up-regulated DEGs was computed using Fisher’s exact test in (i) excitatory neurons across six regions individually and jointly, and (ii) six major brain cell types with regions combined. Analyses were performed under both pathological (NIA-Reagan score) and clinical (cognitive) AD definitions, as defined by the original study. Pairwise differences between log odds ratios were evaluated using a Z-test based on the standard errors of the log odds ratios.

To validate the functional relevance of the ADNeuronNet interactors, we performed cell type-specific and pathway enrichment analyses. For cell type enrichment, we used the Human Protein Atlas (HPA) database to assess tissue and cell type specificity of our detected interactors. For pathway enrichment, we performed KEGG pathway analysis downloaded from API (https://www.kegg.jp/kegg-bin/show_brite?hsa00001.keg). Enrichment significance was calculated using hypergeometric tests with Benjamini-Hochberg correction (adjusted *p* < 0.05).

To assess the quality of our interactome against background noise, we compared the connectivity of our interactors to a randomized protein-protein interaction (PPI) network. We obtained the human PPI network and generated 1,000 randomized networks by degree-preserving shuffling. For each AD pathology, we calculated the odds ratio of interactions in our observed network compared to the distribution of odds ratios in randomized networks. Statistical significance was assessed using permutation tests, with *p* calculated as the proportion of randomized networks with odds ratios equal to or greater than the observed value.

### AlphaFold3 Structural Modeling

AlphaFold3 (AF3) was used to generate atomic-resolution structural models for all bait-interactor pairs in ADNeuronNet, and has successfully generated structures for 1,570 interactions, excluding interactions involving APOE variants. For benchmarking, AF3 was applied to protein pairs derived from 2,075 experimentally resolved PDB co-complex structures, including 562 interacting (positive) and 3,650 non-interacting (negative) pairs. We additionally modeled 540 interactions from the OpenCell dataset (128 interactions involving 26 shared baits with ADNeuronNet and 412 randomly selected) and 4,580 randomly selected protein pairs without prior evidence of interaction and not found in the same PDB co-complex as further controls. All predictions were generated using default settings, and interaction confidence was evaluated using standard AF3 interface metrics.

### N2A Immunofluorescence Imaging

Mouse neuroblastoma N2A cells were maintained in DMEM (Gibco) supplemented with 10% fetal bovine serum. BIN1 isoform 1 cDNA (a kind gift from Dr. Raja Bhattacharyya, Massachusetts General Hospital, Harvard Medical School) was subcloned into the FUGW lentiviral vector. N2A cells were co-transfected with FUGW-BIN1v1-GFP and pCMV-mCherry-RAB5 (Addgene #55126), alongside pCMV-Flag-RIN2 or pCMV-Flag-RIN3 (kindly provided by Dr. Hiroaki Kajiho, Tokyo Medical and Dental University), using polyethylenimine (PEI). Following transfection, cells were seeded onto poly-L-lysine-coated coverslips to promote adherence.

Cells were washed with PBS and fixed with 4% paraformaldehyde (PFA) for 15 minutes at room temperature, followed by three PBS washes. Primary antibodies anti-Flag (Sigma-Aldrich, F1804) were diluted 1:200 in Intercept (PBS) Blocking Buffer (LI-COR, 927-70001) containing 0.05% Saponin (Thermo Fisher Scientific, J63209.AK) for permeabilization. After overnight incubation at 4°C, coverslips were washed three times (5 minutes each) with PBST (PBS + 0.1% Tween-20). After washing, coverslips were incubated with Goat Anti-Mouse IgG H&L (Alexa Fluor® 647) (1:200, ab150115, Abcam) for 1 h at room temperature in blocking buffer containing Saponin. Coverslips were washed and mounted onto glass slides using Fluoromount-G™ (Thermo Fisher Scientific, 00-4958-02) and imaged via confocal microscopy using 100X objective. Fluorescently tagged proteins (e.g., BIN1V1-GFP, mCherry-RAB5) were directly imaged, as the labels provided sufficiently strong signals.

### Knockdown iPSC Neuron Preparation

4R-P301S iPSCs were plated at a density of 1.2 × 10⁷ cells per 10-cm dish in pre-differentiation medium. On day 0, pre-differentiated neurons were replated either at 1 × 10⁶ cells per well onto PDL–coated 6-well plates or at 2 × 10⁵ cells per well onto laminin-coated coverslips placed in PDL-coated 24-well plates and maintained in maturation medium.

On day 3, lentiviral particles encoding control shRNA or shRNAs targeting ANAPC1 or ANAPC2 were added to the cultures. After 24 h, virus-containing medium was replaced with fresh maturation medium. Beginning on day 7, iPSC-derived neurons were exposed to 2 μg/mL K18 fibrils for a total duration of 4 weeks. On day 35, neurons cultured in 6-well plates were collected for western blotting and proteomic analyses, while neurons grown on coverslips were fixed for immunocytochemical studies.

### Human iPSC Immunocytochemistry and Imaging

Human iPSCs and iPSC-derived neurons were fixed with 4% paraformaldehyde (Electron Microscopy Sciences) in PBS for 15 min, followed by three washes (5 min each) with DPBS containing Mg²⁺ and Ca²⁺ (Corning, 21-030-CM). Cells were permeabilized using 0.1% Triton X-100 in DPBS for 10 min and subsequently blocked for 1 h at room temperature in DPBS containing 0.1% Triton X-100 and 5% normal goat serum.

Primary antibodies diluted in blocking buffer were applied overnight at 4°C. After three washes with DPBS, cells were incubated with fluorophore-conjugated secondary antibodies (1:500; Alexa Fluor® 488, 568, or 647; Thermo Fisher Scientific) for 1 h at room temperature. Primary antibodies included chicken anti-MAP2 (Novus Biologicals, NB300-213) and mouse anti-conformationally abnormal Tau (MC1, Peter Davies). Coverslips were washed three additional times and mounted using Vectashield mounting medium containing DAPI (Thermo Fisher Scientific, H-1200-10).

Images were acquired using either a Keyence BZ-X710 microscope or a Zeiss LSM 880 laser scanning confocal microscope and processed with Zen 3.2 software. Imaging settings were adjusted to avoid saturation of the majority of high-intensity pixels. Quantitative analyses were performed using FIJI (NIH).

### Western Blot

To validate the interaction between RIN2 and BIN1, either iPSC-derived neuron lysate (5 × 106 cells) or 350 µg of commercial Human Brain Whole Tissue Lysate (Novus Biologicals) was used. Samples were incubated overnight at 4 °C with Dynabeads™ Protein G (Novus Biologicals, NB820-59177) conjugated to either 2.5 µg of RIN2 antibody or a Rabbit IgG Isotype Control (Thermo Fisher Scientific, 26102). Protein-bound Dynabeads were washed three times with Digitonin lysis buffer, resuspended in SDS protein loading buffer, and heated at 95 °C for 5 min. Proteins were separated on 4–20% SDS-PAGE gels (Bio-Rad, 4561093) and transferred to PVDF membranes. Endogenous RIN2 and BIN1 were detected using PA5-110299 (Thermo Fisher Scientific) and 14647-1-AP (Proteintech) antibodies, respectively.

To validate the interactions of RIN3-SNX9 and RIN3-SNX18, the changed interactions between BIN1 isoforms, and the changed CD2AP interaction between wildtype (WT) RIN3 and RIN3-W63C, each gene variant was cloned into the FUGW vector as previously described. Wildtype RIN3 cDNA used for cloning was kindly gifted by Dr. Hiroaki Kajiho. Site directed mutagenesis was carried out using the following primers to generate RIN3-W63C (Forward 5’-CTCATCAAAACATGCCCGGTGTGTCTGCAGCTGAGTCTGGGCCAGG-3’ and 5’ CCTGGCCCAGACTCAGCTGCAGACACACCGGGCATGTTTTGATGAG-3’). Human iPSC-derived neurons expressing BIN1v1-GFP, BIN1v9-GFP, RIN3-GFP, RIN3 W63C-GFP, or insert-free FUGW vector, as well as uninfected i3Neurons were generated and immunoprecipitated with anti-GFP beads as described in previous sections. Proteins were separated with 4-20% SDS-PAGE gels (Bio-Rad, 4561093) and transferred on PVDF membranes (Cytiva, 10600023). Membranes were blocked with 5% milk in TBST (TBS + 0.1% Tween-20) or PBST (PBS + 0.1% Tween-20), then incubated with primary and secondary antibodies diluted in TBST or PBST. Primary antibodies included anti-SNX9 (Proteintech,15721-1-AP), anti-SNX18 (Proteintech, 21946-1-AP), anti-GFP (Genetex, GTX 628528), anti-AP2A1 (Proteintech, 29887-1-AP), anti-AP2B1 (Proteintech, 15690-1-AP), anti-TOMM40 (Proteintech, 18409-1-AP), and anti-CD2AP (Proteintech, 51046-1-AP). Secondary antibodies for signal detection were Donkey anti-Rabbit IgG (Alexa Fluor® 680) (Invitrogen, A10043) and Donkey anti-Mouse IgG (Alexa Fluor® 680) (Invitrogen, A10038). ImageJ (NIH) was used to quantify densitometry of the bands. For densitometric analysis, band intensities were first background-corrected by normalizing each target band to the corresponding signal in the insert-free FUGW control lane. To account for differences in RIN3 abundance and pulldown efficiency between WT and W63C samples, a scaling factor was derived by equalizing the normalized RIN3 band intensities across the two conditions. This scaling factor was subsequently applied to the normalized CD2AP signal in the RIN3-W63C samples. The adjusted CD2AP intensities were then used for quantitative comparison and visualization. Statistical analyses were performed using a one-tailed Student’s t-test to evaluate directional differences between WT and RIN3-W63C conditions. The t statistic and corresponding *p* is reported in the figure.

To perform co-IPs of BIN1-ANAPC1 and BIN1-ANAPC2 and validate protein expression perturbed in the shRNA knockdown cell lines, human iPSC-derived neurons were washed twice with ice-cold DPBS (Thermo Fisher Scientific, 14190144), pelleted by centrifugation at 300 × g for 5 min at 4°C, and lysed in cold RIPA buffer (Thermo Fisher Scientific) supplemented with protease inhibitors (Sigma-Aldrich, P8340), phosphatase inhibitors (Sigma-Aldrich, P5726 and P0044), and deacetylase inhibitors, including nicotinamide (Sigma-Aldrich, 72340) and trichostatin A (Sigma-Aldrich, T8552). Lysates were incubated on ice for 10 min and clarified by centrifugation at 21,000 × g for 10 min at 4°C. Supernatants were collected, and protein concentrations were measured using the Pierce BCA Protein Assay Kit (Thermo Fisher Scientific). Equal amounts of protein were separated on 4–12% SDS–PAGE gels (Invitrogen) and transferred onto nitrocellulose membranes (Bio-Rad, 1620115). Membranes were blocked with 5% milk in TBST and incubated with primary and secondary antibodies diluted in blocking buffer. Primary antibodies included anti-ANAPC1 (Proteintech, 21748-1-AP), anti-ANAPC2 (Proteintech, 13559-1-AP), anti-APOE (Proteintech, 84775-6-RR), anti-TOMM34 (Abclonal, A27418), and anti-BIN1 (Sigma-Aldrich, 05-449). HRP-conjugated secondary antibodies (Sigma-Aldrich) and chemiluminescent substrates (Bio-Rad) were used for signal detection. Densitometric quantification of immunoblot bands was performed using Image Lab (Bio-Rad) or FIJI (NIH).

To validate interactions in AD patient-derived neurons, mature neurons were lysed using NP-40 lysis buffer (10 mM Tris-HCl, pH7.4, 120 mM NaCl, 2 mM EDTA, 0.5% NP40) with protease and phosphatase inhibitors (Thermo Fisher Scientific). The protein concentration of lysates was quantitated using Pierce BCA assay (Thermo Fisher Scientific). 250 µg proteins and 1 µg indicated antibody (the corresponding IgG as control antibody) were mixed and rotated overnight at 4 °C. 10 µl Dynabeads Protein G were then added into the antibody-protein mixture for 30 min at 4 °C. The beads were washed with 500 µl IP buffer twice. The protein complex captured by the indicated antibody was eluted with SDS elution buffer and subjected to western blot. Signals were detected using Clarity ECL substrate kit (Bio-Rad) on Amersham imager 600. Primary antibodies included anti-PICALM (Proteintech, 28554-1-AP), anti-CD2AP (Proteintech, 51046-1-AP), anti-AP2B1 (Proteintech, 15690-1-AP), anti-CLTC (Proteintech, 26523-1-AP), anti-CAPZB (Proteintech, 25043-1-AP), and anti-CAPZA2 (Proteintech, 15948-1-AP).

## Data Availability

The raw mass spectrometry data of all IPs have been deposited to the ProteomeXchange Consortium via the PRIDE^101^ partner repository with the dataset identifier PXD075357.

## Code Availability

Jupyter notebooks, python and R scripts, and associated datasets, used for analysis and figure generation are available on GitHub (https://github.com/haiyuan-yu-lab/ADNeuronNet).

## Author Contributions

H.Y. and X.W. conceived and oversaw all aspects of neuron interactome mapping; Yuansong W. and L.G. advised on all aspects of iPSC culture/differentiation and functional interpretation/validation; H.Y., F.C., and L.G. designed all analyses; Interactome mapping: X.W., K.M, Y.S., Yiwen W., and Y.T.; Analyses: Y.S., T.X., Y.H., Y.K., Z.S., and Y.Z.; RIN2 functional study: X.W. with input from Yuangsong W. and W.L.; APC/C-APOE functional study: Yuansong W. led the effort and worked with K.M., Y.S., W.F., S.L., and J.Z; co-IPs: X.W., K.M., Yuansong W., and W. S.

## Acknowledgements

We used instruments supported by the BRC Flow Cytometry Facility (RRID:SCR_021740) of the Biotechnology Resource Center of Cornell Institute of Biotechnology, and thank them for their assistance with our FACS work. We thank Yilin Liu for assistance with AlphaFold3 structural modeling, Dr. Min Wan in Dr. Yuxin Mao’s lab for assistance with confocal imaging, Dr. Hiroaki Kajiho from Tokyo Medical and Dental University for providing RIN2 and RIN3 constructs, and Dr. Raja Bhattacharyya from Harvard Medical School for providing BIN1v1 cDNA. This work was supported by grants from the National Institute of Aging (NIA): R01 AG092462 to H.Y., L.G., and F.C, RF1 AG082211 to H.Y. and F.C, R01 AG077899 to H.Y. and L.G., U01 AG073323, R01 AG066707, R01 AG084250, R01 AG076448, R01 AG082118, R01AG092591, and R33AG083003 to F.C., as well as grants from National Institute of Neurological Disorders and Stroke (NINDS, RF1 NS133812, the Alzheimer’s Association award (ALZDISCOVERY-1051936), and the Alzheimer’s Disease Drug Discovery Foundation (ADDF) to F.C.

## Notes

### Competing Interest Statement

The authors have declared no competing interest.

