## Supplementary Figures for "Interactome mapping in human excitatory neurons reveals novel risk genes and pathways in Alzheimer’s disease"

**Supplementary Figure S1. Protein length distribution of AD risk proteins and overlap of BIN1v9 interactors across lentiviral infection and cell-sorting conditions.**
**(a)** Distribution of protein length (amino acids) for 97 neuron-expressed AD risk proteins, stratified by cloning outcomes.
**(b)** Infection of iPSCs followed by fluorescence-activated cell sorting (FACS) of GFP-positive cells is necessary to obtain sufficient material for IP-MS interactome mapping. Interactors were identified as the proteins enriched in the BIN1v9 IP samples passing cutoffs of log2FC ≥2 and adjusted p-value <0.05. We performed DDA-TMT quantification during this stage of the optimization process. Scale bar on confocal microscopy images represents 100μm.

**Supplementary Figure S2. Comparison of DDA and DIA MS methods.** A spike-in benchmarking dataset consisting of yeast proteome spiked into human proteome at different ratios was used to test the performance of DDA-TMT vs DIA-LFQ methods. In this model, yeast proteins always have significant changes across different samples (thus, could be considered as representation of true interactors in an IP-MS experiment), while human proteins remain unchanged across samples (representing non-specific binders in IP-MS experiments). Compared to the DDA-TMT workflow, the DIA-LFQ workflow achieves higher proteome coverage (>11k proteins identified vs. ~4k) with better quantitative performance (~98% successful rate vs ~92%).

**Supplementary Figure S3. ADNeuronNet interactors enrichment for AD up-regulated genes across brain regions, cell types, and diagnostic definitions**
**(a)** Enrichment of up-regulated genes in pathologically diagnosed AD dementia patients in excitatory neurons. Differentially expressed genes were derived from snRNA-seq analysis of ROSMAP samples (26 AD, 22 non-AD). Enrichment was calculated as log2 fold-change using Fisher's exact test and compared to interactors in literature protein interactions and combined BioPlex & OpenCell interactomes. Pairwise differences were assessed using Z-test based on the standard errors of the log odds ratios.
**(b)** Cell-type–specific enrichment of AD up-regulated genes across all brain regions combined using clinical (cognitive) diagnosis. Enrichment and statistical comparisons were performed as in (a).
**(c)** Cell-type-specific enrichment of AD up-regulated genes across all brain regions combined using pathological diagnosis. Enrichment and statistical comparisons were performed as in (a).

**Supplementary Figure S4. Distribution of AlphaFold3 ranking score values across different interaction datasets indicates high quality of ADNeuronNet interactions.**

**Supplementary Figure S5. Co-immunoprecipitation western blot validation for known interactions in human WTC11 derived neurons and AD patient iPSC derived neurons.**
**(a)** RIN3 interacts with SNX18 in human WTC11 derived neurons.
**(b-c)** PICALM interacts with AP2B1 and CLTC in **(b)** human WTC11 derived neurons and **(c)** AD patient iPSC derived neurons.
**(d-e)** CD2AP interacts with CAPZB and CAPZA2 in **(d)** human WTC11 derived neurons and **(e)** AD patient iPSC derived neurons.

**Supplementary Figure S6. Determination of changes in interaction strength between wild-type (WT) and mutant conditions and BIN1 interaction networks from previous large-scale human interactomes.**
**(a)** The WT and mutant IPs are first compared to the GFP control IPs to determine the high-confidence interactors of each. Then, the abundance values for the union of identified interactors are compared between the two conditions to determine if the interaction was strengthened, gained, weakened, or lost.
**(b)** BIN1 interactome generated in HEK293T cells from OpenCell (Cho et al., 2022).
**(c)** BIN1 interactome generated in HEK293T and HCT116 cells from BioPlex (Huttlin et al., 2021).

**Supplementary Figure S7. Functional consequences of depleting BIN1 interacting APC/C components ANAPC1 and ANAPC2.**

**(a)** Volcano plot of quantitative proteomics showing differentially expressed proteins in human 4R neurons transduced with shCtrl or shANAPC2 #2, with APOE identified as the top upregulated protein following ANAPC2 knockdown.

**(b–c)** Venn diagrams illustrating overlap among upregulated **(b)** and downregulated **(c)** differentially expressed proteins induced by shANAPC1 and by two independent shANAPC2 constructs (#1 and #2).

**(d–e)** Pathway enrichment analyses of shared upregulated **(d)** and downregulated **(e)** differentially expressed proteins following ANAPC1 and ANAPC2 depletion.

**(f–h)** Immunoblot analysis and quantification of ANAPC1 and APOE protein levels in human neurons transduced with lentiviral shANAPC1 alone or in combination with shAPOE, confirming efficient knockdown and modulation of APOE expression.

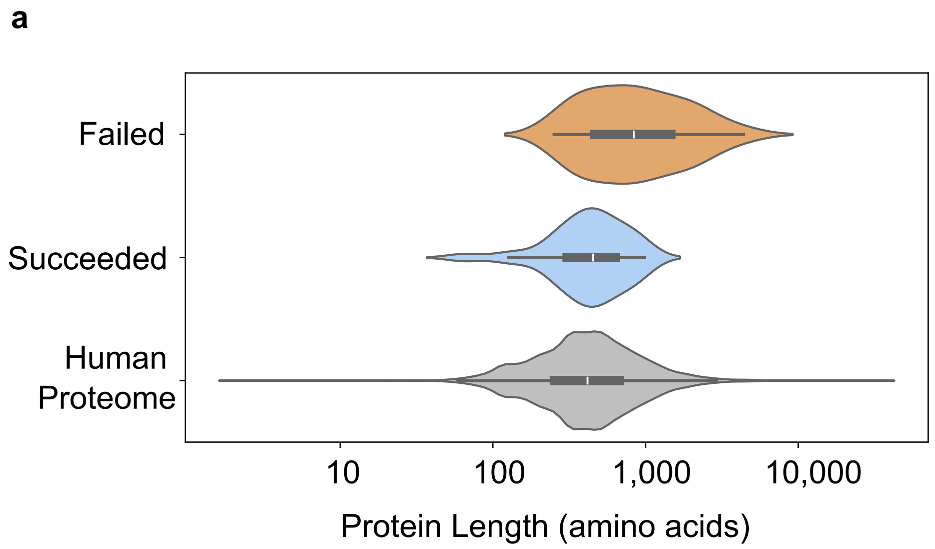


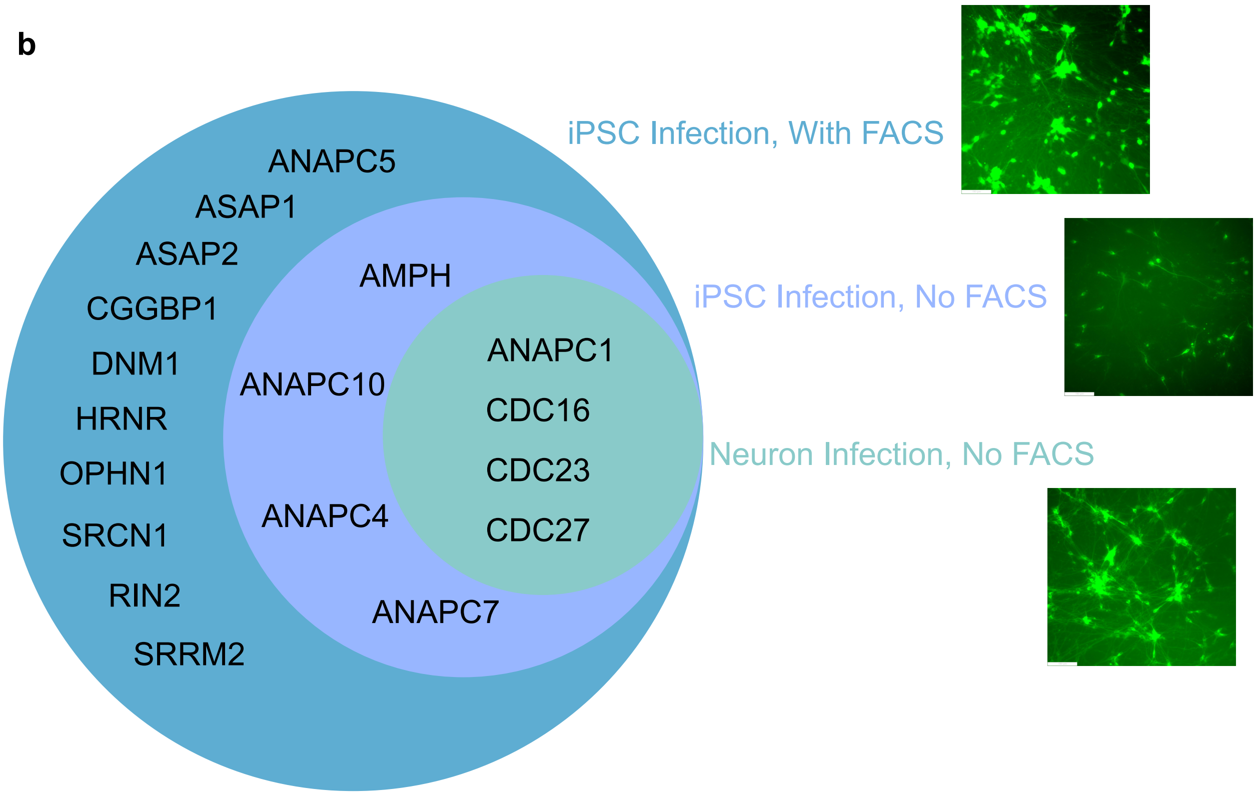


#### Supplementary Figure S1

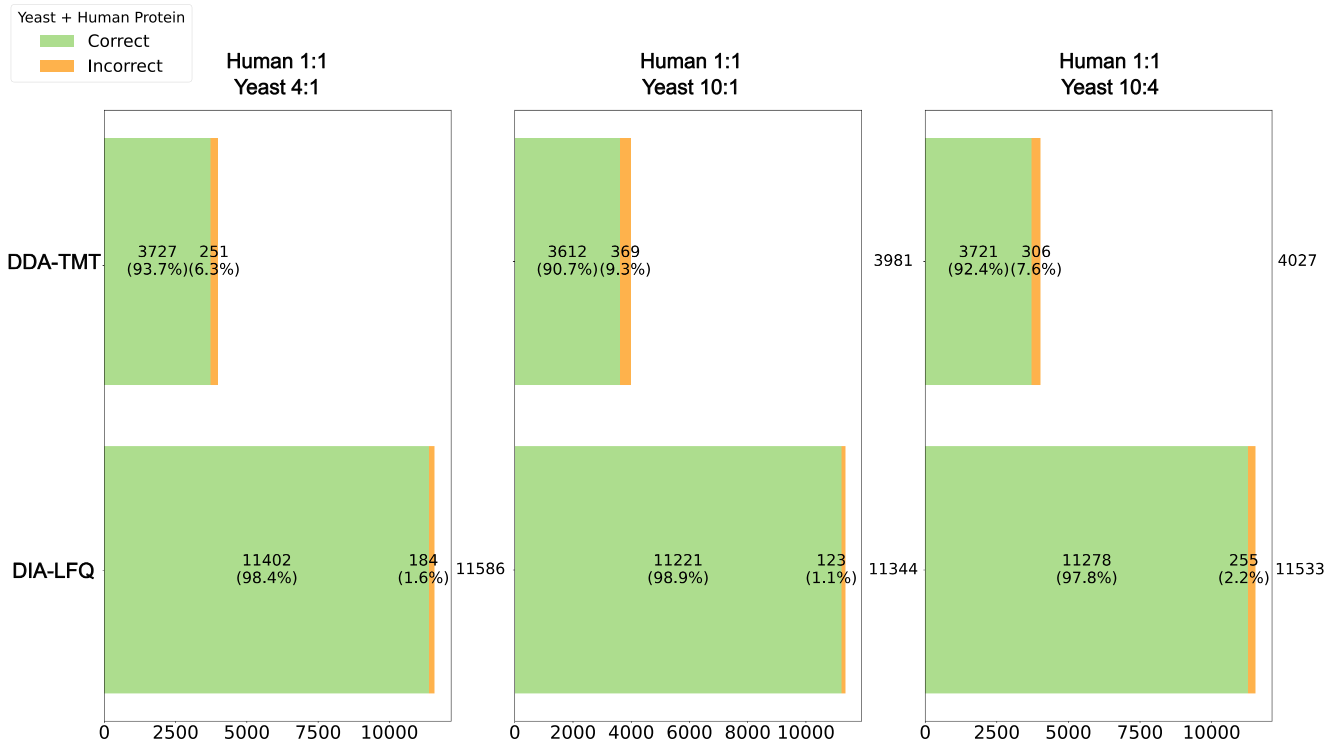


### Supplementary Figure S2

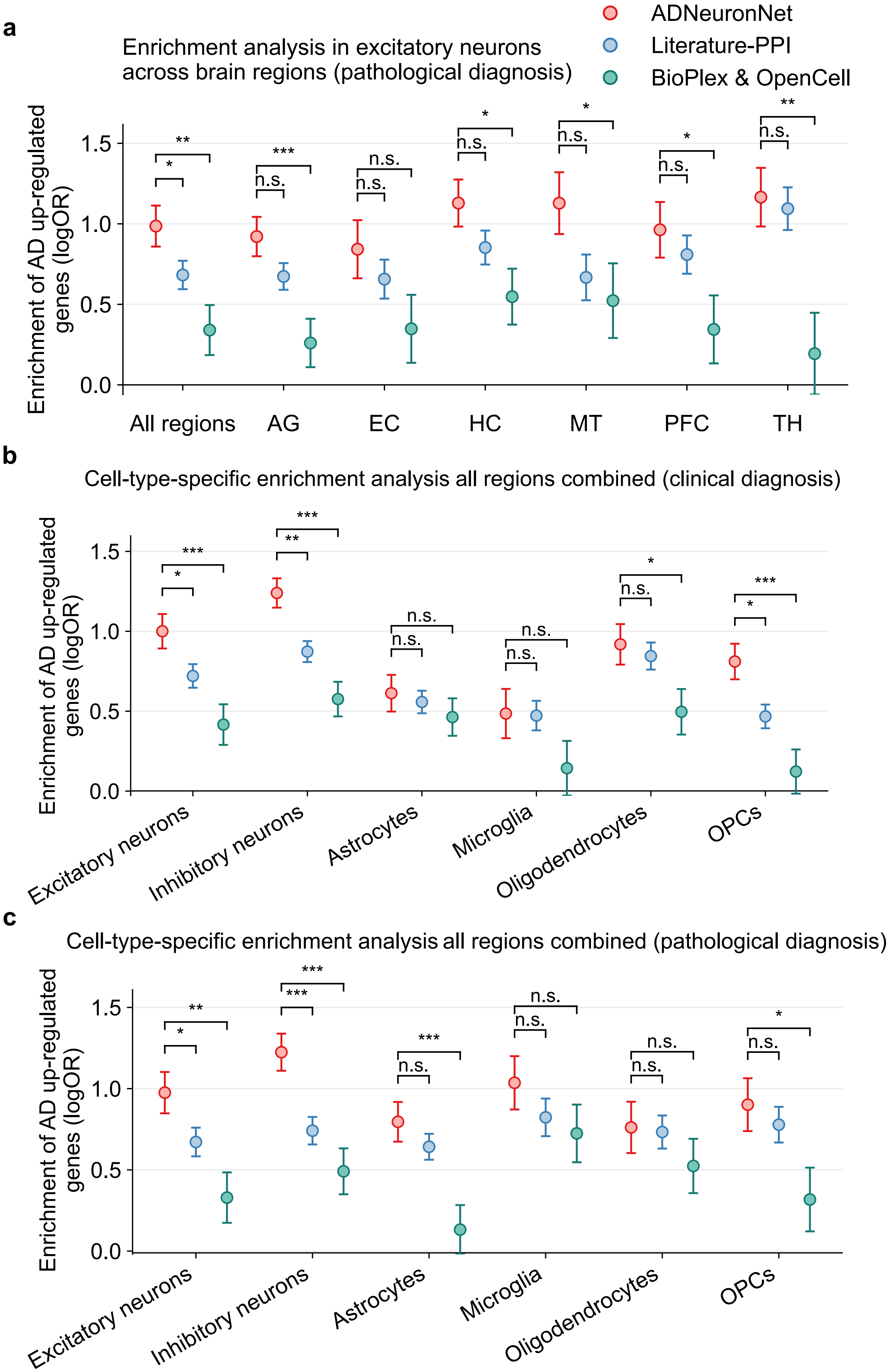


**Supplementary Figure S3**

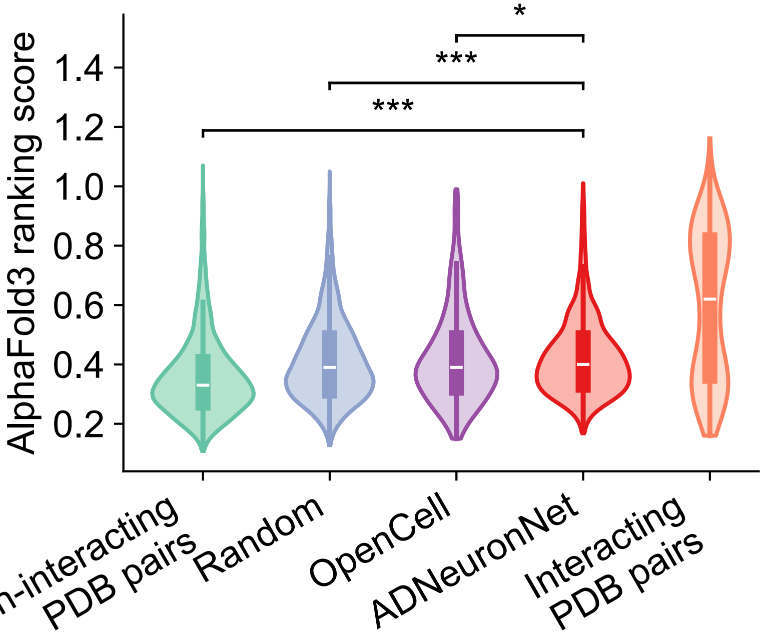


**Supplementary Figure S4**

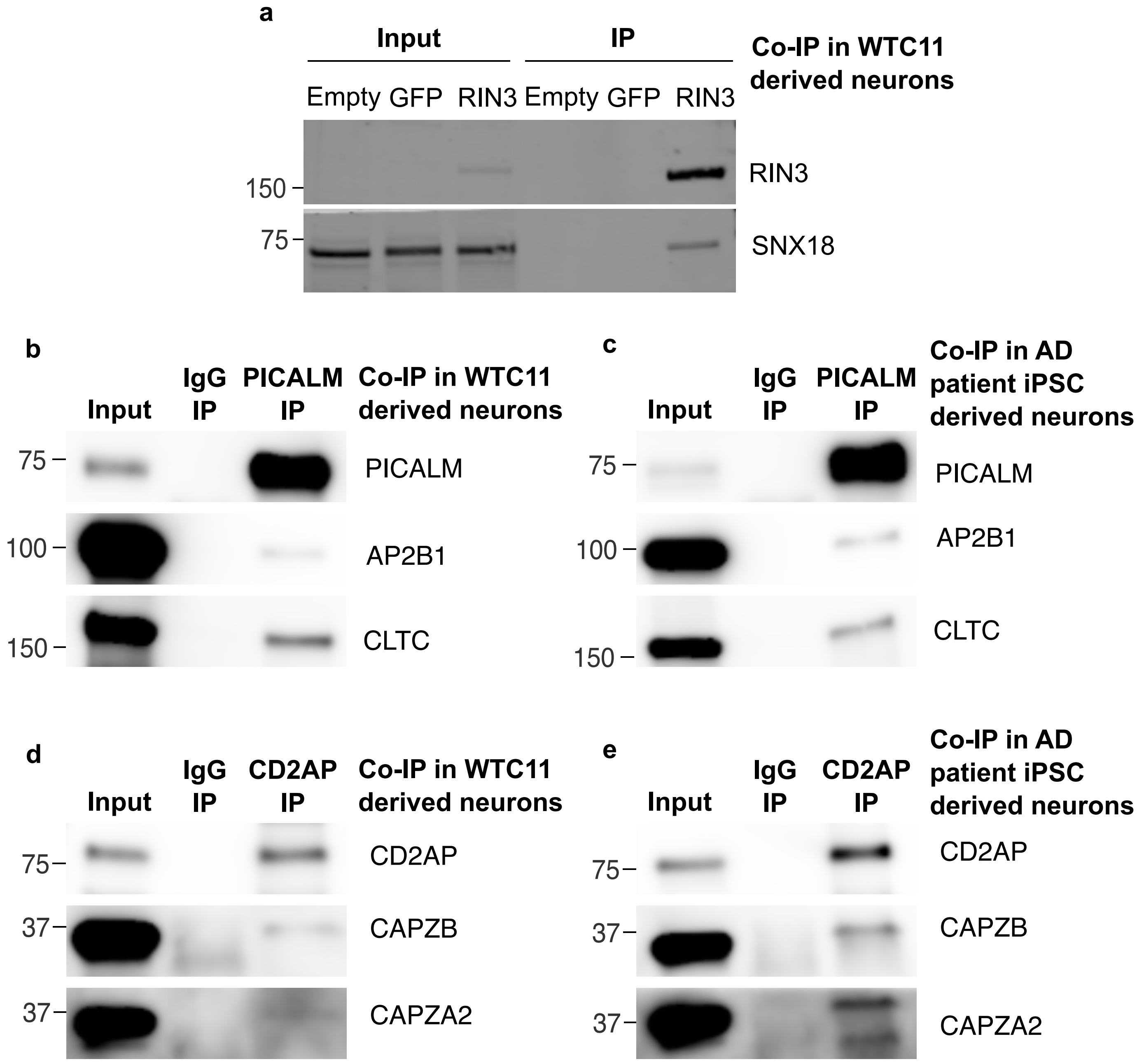


### Supplementary Figure S5

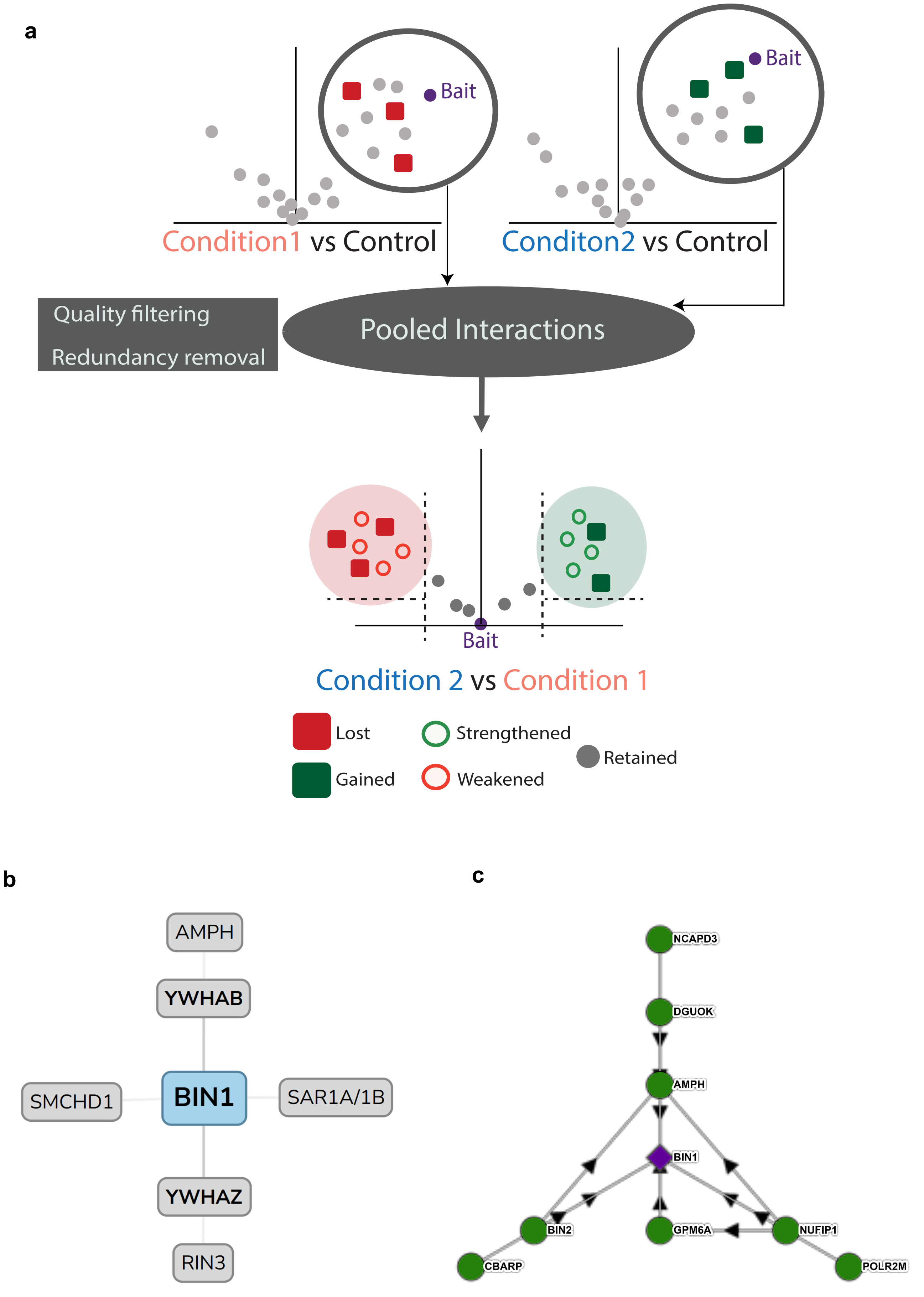


**Supplementary Figure S6**

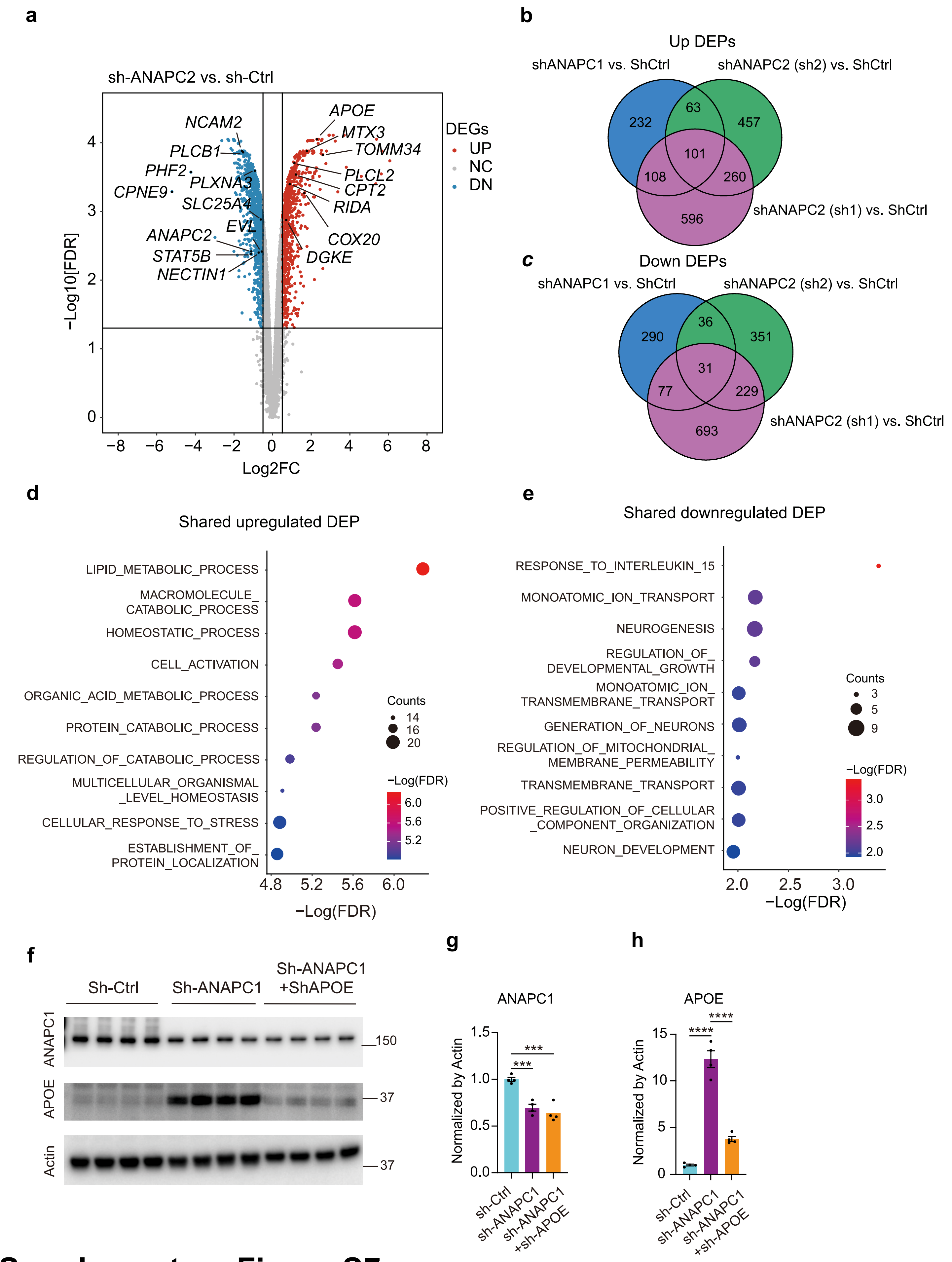


### Supplementary Figure S7
